# How Cerebellar Architecture and Dense Activation Patterns Facilitate Online Learning in Dynamic Tasks

**DOI:** 10.1101/2022.10.20.512268

**Authors:** Adriana Perez Rotondo, Dhruva V. Raman, Timothy O’Leary

## Abstract

The cerebellum has a distinctive architecture in which inputs undergo a massive size expansion in dimensionality in early layers. Marr and Albus’s classic codon theory and more recent extensions^1–4^ argue that this architecture facilitates learning via pattern separation. The essence of this idea is that sparsely active clusters —‘codons’— of inputs are more easily separable in a higher dimensional representation. However, recent physiological data indicate that cerebellar activity is not sparse in the way anticipated by codon theory. Moreover, there is a conceptual gap between static pattern separation and the critical role of the cerebellum in dynamic tasks such as motor learning. We use mathematical analysis and simulations of cerebellar learning to identify specific difficulties inherent to online learning of dynamic tasks. We find that size expansions directly mitigate these difficulties, and that this benefit is maximised when granule cell activity is dense.

## Introduction

The cerebellar cortex has a distinct circuit architecture characterised by a large ‘input expansion’: each mossy fibre input projects to approximately 250 granule cells, the wider population of which comprises more than half of the neurons in the entire brain.^5^ Speculation on the purpose of this costly size expansion has spurred a long line of scientific studies over the last half century.^6^

One of the earliest and influential proposals is the codon theory of Marr and Albus.^1,2^ This theory and its more recent refinements have shown how size expansions can facilitate pattern separation.^4,7,8^ Pattern separation in this context means that distinct signals embedded in overlapping cerebellar inputs can be accurately resolved into less correlated cerebellar output signals. This may enable refined behavioural responses to complex features in the animal’s internal and external state, and thus aid learning in a variety of settings.

A key prediction of Marr-Albus theory is that the granule cell layer activity should be sparse, with only around 5 − 10% of granule cells active at a time^1,2,9–11^ Yet, recent experimental studies have found dense granule cell layer activity in motor control tasks,^12–16^ throwing this theory into doubt. Furthermore, there is a conceptual gap between the proposed functional role of pattern separation, and the actual functional roles that the cerebellum is known to be involved in. We know that cerebellar function is necessary for the precise execution and real-time calibration of movements,^17–21^ but it remains unclear how pattern separation aids in this computational task.^11^

We bridge these gaps between theory and observation by developing an alternative analysis that explains how the cerebellar input expansion specifically impacts learning in a dynamic, online setting. Using a canonical motor learning framework first proposed by Kawato et al.,^22^ we show that learning errors arise from the requirement to learn from a finite time window during continual execution of a motor task. These errors are due to a bias in estimating gradients in the online setting that will impede any local learning rule. The analysis is therefore independent of the specific plasticity mechanisms operating in this circuit. Our analysis then shows that the impact of this online error is mitigated as additional, apparently redundant granule cells are introduced. All of our theoretical results are validated in a computational model.

Our findings suggest that cerebellar input expansions facilitate learning during the ongoing execution of dynamic tasks by specifically mitigating biases due to online error gradients. This provides an alternative to the Marr-Albus account of one of the cerebellum’s most striking and metabolically costly architectural features. Furthermore, in contrast to Marr-Albus codon theory, we find that this expansion is more effective when a sufficient number of units in the expanded representation are active. Optimal learning conditions within our model therefore predict the dense granule cell activity that has been observed in cerebellar cortex recently.^12–16^

## Results

### A standard model of cerebellar learning

Throughout this study we refer to a canonical model of motor control, due to Kawato.^22,23^ We will briefly recapitulate the model then use it to motivate the results in the remainder of the paper and validate theoretical analyses.

Figure 1A is an outline of the structure of the model, in which the motor cortex and cerebellum cooperate to produce precise motor output trajectories. The model assumes that input signals, corresponding to desired movements, are transformed into motor commands that are issued to the musculoskeletal system, sometimes called the motor plant. These resulting motor commands are modified continually via negative feedback from sensory signals that convey mismatch between the desired and actual trajectories (see Methods).^24^

**Figure 1:**
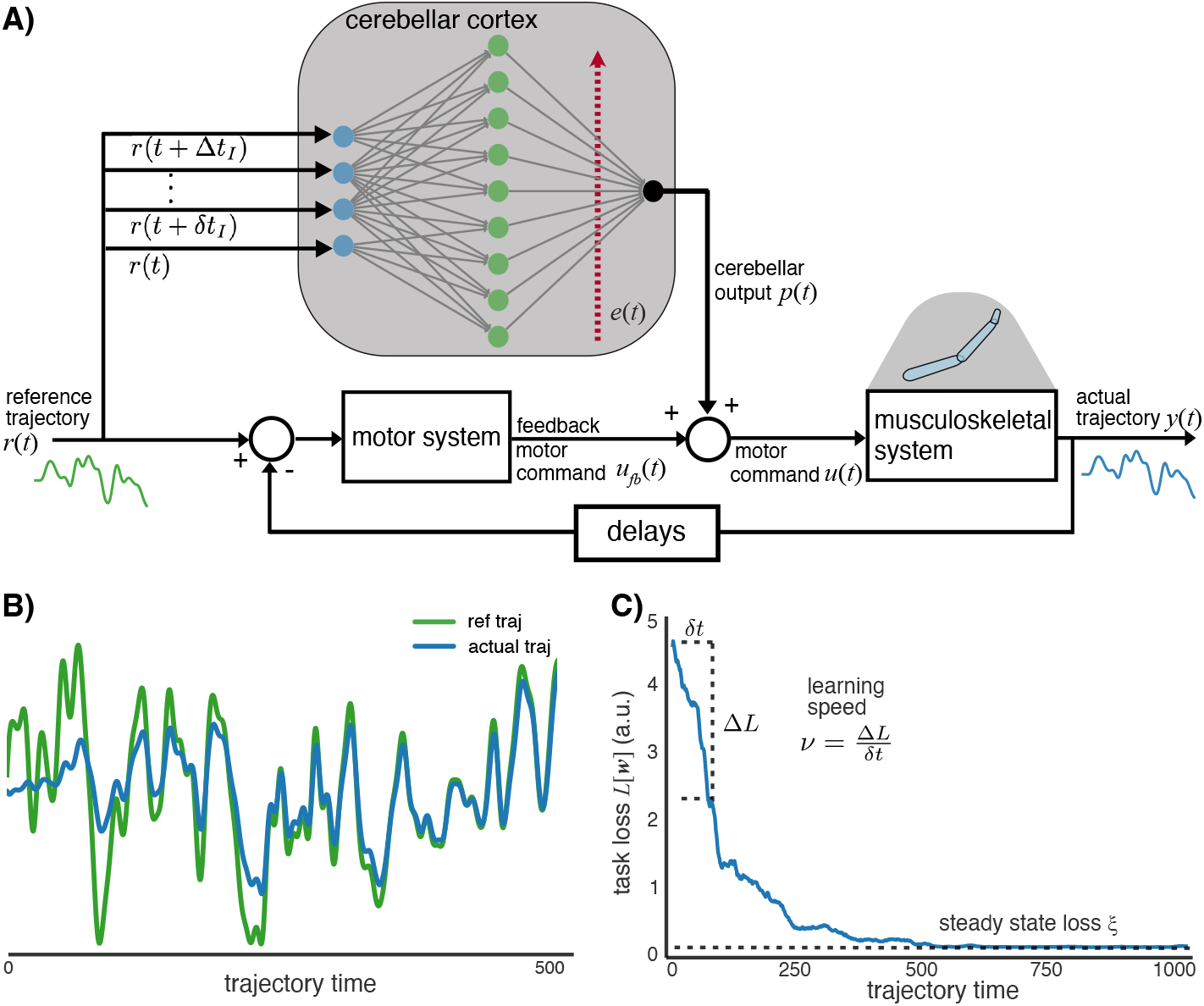
System used to model the cerebellum for motor control and learning. **A)** Diagram of the system for the trajectory tracking task. The goal of the task is for the motor plant, the musculoskeletal system, to produce the reference trajectory. The feedback controller represents the part of the motor system which generates an approximate motor command. The cerebellar network transforms a motor plan into a modulatory motor command. The sum of the two motor commands constitutes the total command sent to the motor plant. **B)** The cerebellar cortex network learns online, during the movement, by modifying the output weights according to the error information carried by the climbing fibres. We choose an LMS inspired learning rule given by eq (5). Initially the actual trajectory is far form the reference trajectory. As learning progresses, the two curves come closer together. **C)** The task loss *L*[**w**] defined in eq (6) quantifies the error between the reference and actual trajectories over the whole trajectory, if the weights were fixed to some value **w**. We calculate the task loss after each weight update. The task loss decreases throughout the movement until it reaches a steady state value referred as the *steady state loss*. The *learning speed* quantifies how fast the task loss decreases initially.

By itself, negative feedback cannot perfectly correct for mismatches between a planned and desired trajectories in all circumstances. Biophysical delays between a movement mismatch, its sensation, and its correction will result in errors and such delays can even destabilise the system entirely if the feedback gain is too high.^25–29^

The role of the cerebellum in this model is to supplement motor commands with a learned, predictive signal that compensates for the limitations of negative feedback.^30^ Conceptually, the cerebellum thus constitutes an *inverse model* of the motor system: a prediction of which motor commands are required to realise a particular output trajectory.^31^ This inverse model allows the cerebellum to refine approximate motor commands sent by other brain areas to produce a desired movement.^32–37^ Because desired movements, along the properties of the environment and motor plant are all subject to change, the cerebellum must continually update this model.

We model the cerebellum as receiving a desired trajectory, *r*(*t*), extending into the future to some finite horizon, Δ*t_I_*. Anatomically, this signal is believed to reside in mossy fibre inputs. For convenience, and to reflect the finite bandwidth of the input signal, we assume it has a finite sampling resolution *δt_I_*. Thus, the vector of all the *I* mossy fibre firing rates at time *t* is given (Figure 1A as

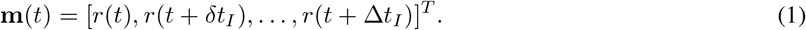

The input is first projected to the cerebellar granule layer, consisting of *N* granule cells. We model the firing rate *h_i_*(*t*) of the *i^th^* granule cell as

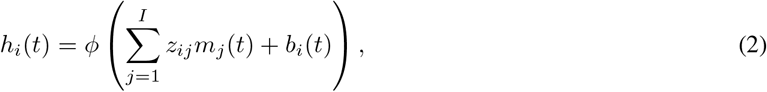

where *ϕ* is the nonlinear activation function, and *z_ij_* is the synaptic weight strength from the *j*’th input mossy fibre to the *i*’th granule cell. The actual firing rate of a granule cell, given a degree of synaptic input from the mossy fibres, depends upon unmodelled factors such as inhibition from basket and stellate cells. Such factors are lumped in the *bias* term *b_i_*(*t*). Granule cell activity then determines the firing rate of a single output Purkinje cell:

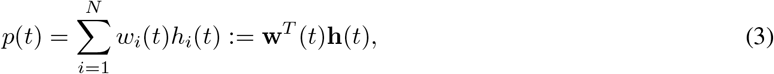

where **w**(*t*) is the vector of synaptic weights carrying signals from granule cells to the Purkinje cell. Anatomically, this corresponds to the parallel fibres/Purkinje cell synapses.

Plasticity in the parallel fibre synaptic weights **w**(*t*) arises from a real-time mismatch between the planned movement and sensory information on the actual movement, and acts to reduce mismatch on future instantiations of the movement.^38–42^ Biologically, this plasticity is induced by complex spikes in the climbing fibres, which activate in a narrow time window (50-200ms) following movement errors.^43–46^

We model this signal as the mismatch, integrated over a narrow time window Δ*t_e_* that terminates by an amount Δ*t_r_* in the past. This temporal offset reflects signal delays in sensory circuitry:

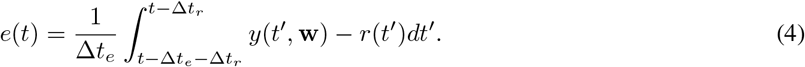

We do not attempt to model in detail the plasticity mechanisms underlying cerebellar adaptation, many of which are the subject of ongoing research. Our goal is to capture general relationships between circuit architecture and learning performance that should hold regardless of the physiological details. Our previous work has shown that we can model any incremental learning rules, regardless of physiological mechanism,^47^ as noise-corrupted gradient descent. We therefore model plasticity as a combination of a gradient term and an uncorrelated, random component:

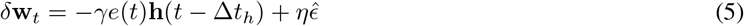

where 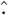 denotes a normalised vector, 0 < *γ* ≪ 1 is the degree of plasticity in the gradient descent direction, Δ*t_h_* is the delay between Purkinje cell input and climbing fibre signal. *ϵ* is a Gaussian vector noise term, scaled by 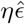 to set the quality of the learning rule, which we sometimes quantify by the signal-to-noise ratio 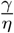. The gradient portion of this learning rule equates to the well-known LMS rule.^48^

Because learning occurs online, synaptic weights in the cerebellar circuit change while the movement is being executed. We assumed that synaptic plasticity operates at a slower timescale (controlled by *γ*) than the dynamics of the firing rates in the system. This is reasonable because synaptic plasticity is slower than firing rate dynamics. This assumption also avoids interference between the two processes that can lead to instabilities (Methods).

### Motivating example

We simulated the Kawato model to quantify how the speed and precision of learning depends on the number of granule cells. We considered an arbitrary trajectory performed over a timespan [0, *T*]. Synaptic weights in the cerebellar circuit, which are randomly initialised (see Methods), update according to the learning rule of (5).

Initially, only negative feedback modulating motor commands from the motor cortex ties the output trajectory to the desired trajectory, resulting in high degrees of mismatch. Over time, cerebellar learning decreases the mismatch, by effectively learning a predictive model of appropriate motor command modulations in response to sensed mismatch. This is visible in Figure 1B.

Because of the cerebellum’s role in facilitating dynamic learning, we next considered how learning performance in this system might be optimised. Naively, one could increase the plasticity induced by a given error signal to speed up learning (increasing *γ* in 5). We refer to this as increasing the *gain* of the learning rule. An excessive gain can, however, slow or destabilise learning: imperfections in the plasticity direction are amplified. This is easy to illustrate by examining the effect of gain on learning speed and steady state loss.

Learning speed quantifies how fast the loss is reduced at initial parts of learning and steady state loss how good the performance is at the end of learning (see Figure 1C and Methods). Excessive learning gain speeds up initial learning, at the cost of steady state precision. Insufficient gain has the opposite effect (Figure 2B). For these reasons we conducted our subsequent analyses of learning performance assuming the biology is doing the best it can in terms of tuning the gain, allowing us to turn our attention to circuit architecture, which is the main focus of our investigation.

**Figure 2:**
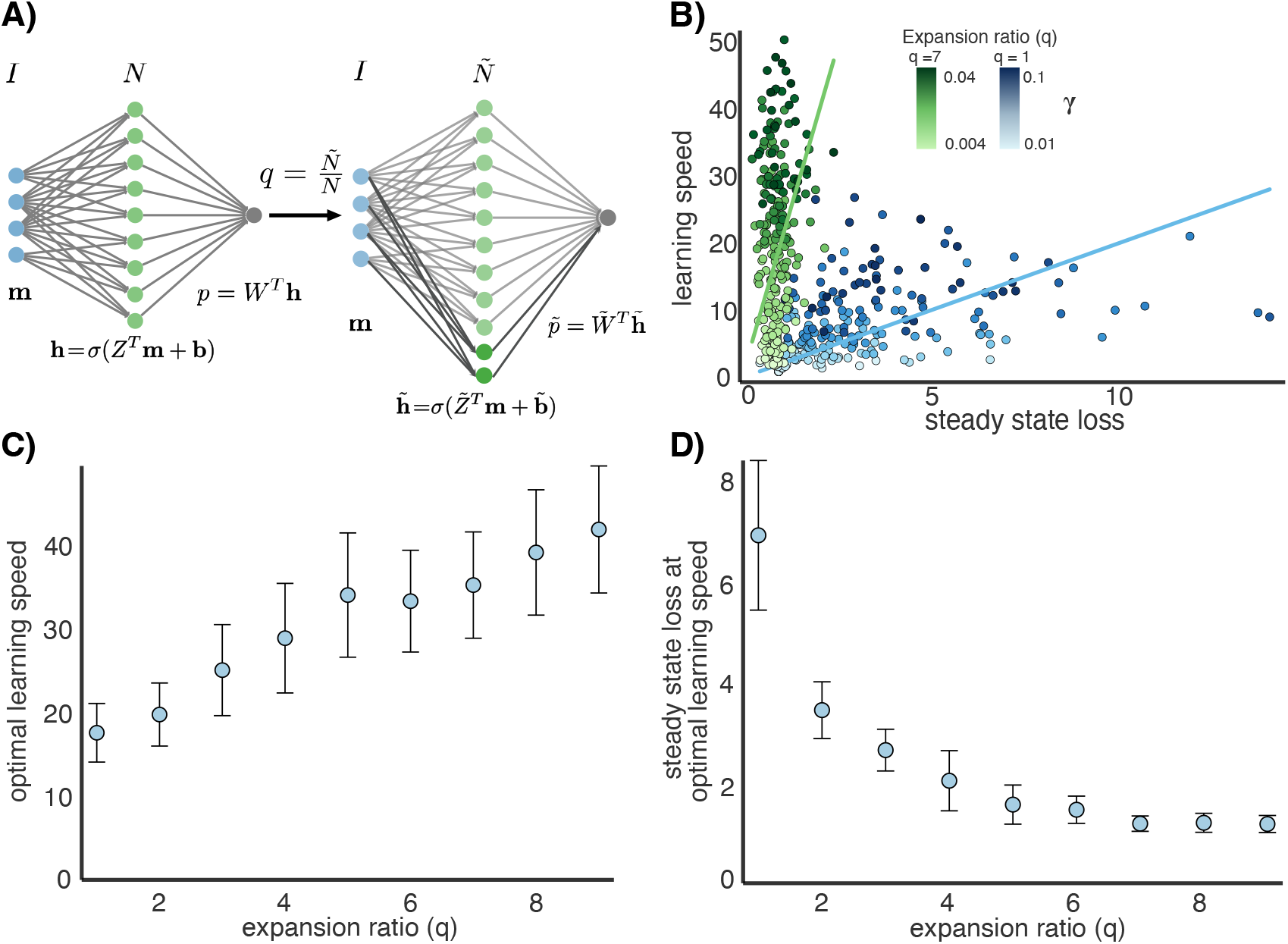
Granule cell layer expansion increases learning performance. **A)** We consider a network expansion that adds granule cells to the hidden layer. The expansion maintains the number of input mossy fibres but increases the number of granule cells from *N* to *Ñ*. The expansion ratio is defined as 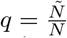. The network expansion changes the granule cell layer activity. **B)** Adding granule cells navigates the trade-off between the learning speed and steady state loss. We plot the trade-off for two different expansion ratios: *q* = 1 in blue, *q* = 7 in green. The shade of the color denotes the value of *γ*. The networks with more granule cells can achieve better learning speed for some steady state loss. **C)** We compute the optimal learning speed over all *γ* for each expansion ratio. The optimal learning speed increases with the input expansion. **D)** We plot the steady state loss for the *γ* that gives optimal learning speed (value in **C)**). The steady state loss decreases with the input expansion indicating that the larger network can achieve better learning speed to steady state loss trade-off. Error bars indicate 1 SEM over simulations with different reference trajectories and weight initialisations. For details on the methods to generate this graph see Methods.

In Figure 2, we successively increase the number of granule cells, while keeping all other aspects of the learning task constant. Each new granule cell forms input connections with *K* = 4 randomly chosen input mossy fibres, and an output connection with the single Purkinje cell (see Methods). Remarkably, both the learning speed and the steady state loss improve, although this improvement saturates (Figure 2C and D). We stress that for both measures we have taken the best achievable values as the plasticity rate, *γ*, is varied. Thus, in this minimal example, ‘excessive’ numbers of granule cells seem to aid learning. In what follows we ask whether this effect is general and unpick its fundamental cause.

### Sources of error in online motor learning

To answer the questions raised in the previous section, we must identify what makes online learning of a trajectory difficult. We need an overall behavioural performance measure across an entire trajectory, which we call the *task loss*:

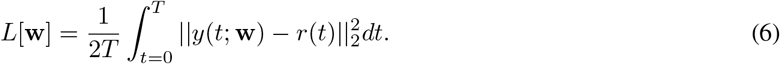

This loss is problematic for the learning system to compute directly because it requires a record of the entire trajectory, and a means of computing the sensitivity across this trajectory to changes in synaptic weight.

Instead, the system uses *online loss* to change weights. This signal is generally believed to reside in climbing fibre activity. We define online loss as:

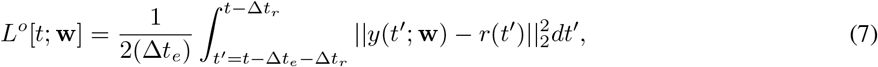

where Δ*t_e_* a time window, and Δ*t_r_* is a delay and *y*(*t*, **w**) is the actual output trajectory. Δ*t_e_* is determined by how much memory of the error the motor system has access to and Δ*t_r_* is determined by climbing fibre delays.^45,49^

As mentioned previously, we will assume that a biologically plausible cerebellar learning rule can only approximate the gradient of the online loss. We therefore model a generic, imperfect, cerebellar learning rule as

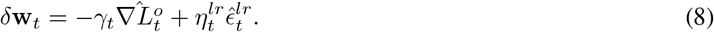

where 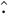 denotes a normalised vector, 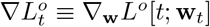 the online gradient. *γ_t_* quantifies the proportion of a weight change in the direction of the online gradient. 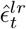 is a unit Gaussian vector. As before, *η^lr^* is a parameter representing quality of the learning rule. We call the term 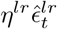 the *learning rule error*. We will refer to learning rules acting according to equation (8), as *online learning rules*.

Systematic differences between online loss and task loss gradients are what make online learning difficult. We next quantify this discrepancy explicitly.

We can rewrite equation (8) in terms of the task loss. First note that

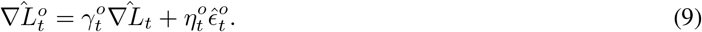

where ∇*L_t_* ≡ ∇_**w**_*L*[**w**_*t*_] is the task gradient. Here, 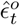 represents the direction of discrepancy between the online and task gradients, and *η°* the magnitude. Together, 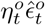 is referred to as the *online error*. Thus, we can rewrite equation (8) as

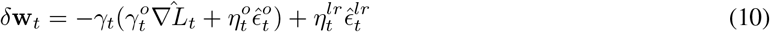

Figure 3B illustrates the decomposition of the change in weights.

**Figure 3:**
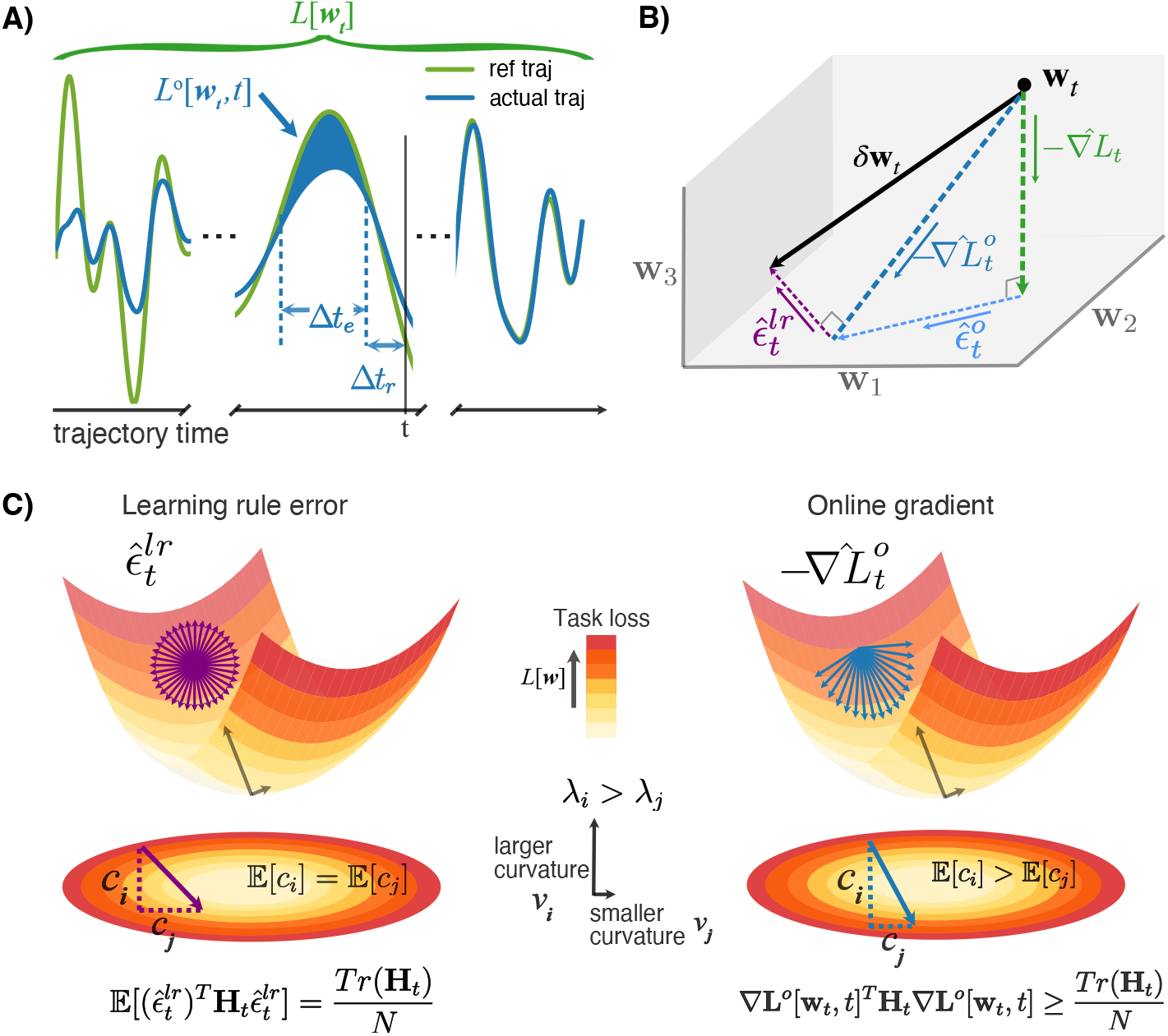
Online learning limits the amount of error information accessible for synaptic plasticity mechanisms to appropriately adjust synaptic weights. **A)** The task loss *L*[**w**] measures the distance between the reference and actual trajectories over the whole movement. At some time *t* during the movement (vertical line), the system only has access to the online loss *L°*[**w**_*t*_, *t*], the distance between the two trajectories over a small window of time in the past (blue shaded region). **B)** The task loss gradient 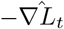 and the online loss gradient 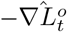 might not be in the same direction in weight space. We refer to the difference between the two as the online error direction 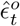. The difference between the direction of change in weights 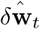 and 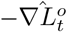 is called the learning rule error direction 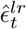. The change in weights can be decomposed as in (10). **C)** Depiction of the projection of the learning rule error (on the left) and the online gradient (on the right) onto the hessian of the task loss. We use a simplified loss function of two weights. On the top, representation of the loss landscape where the z-axis corresponds to the task loss *L*[**w**]. Every point on the landscape corresponds to a different network state **w** with task loss *L*[**w**]. On the bottom, the bird-eye view of the loss landscape. The color represents the value of the task loss. Near a local minimum the loss landscape can be approximated by a quadratic function with constant hessian with two eigenvectors. **v**_*i*_ has a large eigenvalue *λ_i_* and **v**_*j*_ has smaller eigenvalue *λ_j_* on the y and x directions respectively. The learning rule error 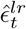 is uncorrelated with the task loss. It can be modelled as a random vector that could be in any direction in weight space (top diagram). On expectation, 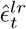 projects evenly onto all the eigenvectors of the hessian (bottom diagram). Hence, on expectation its projection onto the hessian is equal to the average curvature. The online gradient, is not independent of the task loss. It projects more strongly onto directions of larger curvature. Hence its projection onto the hessian will be larger than the average curvature.

### The geometry of the loss landscape controls learning performance

Consider a change in weights *δ***w**_*t*_ over the interval *δt*. From equation (10), this leads to a change in task loss

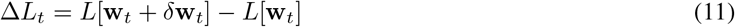

This change in weights results in learning if and only if Δ*L_t_* is negative. Using a Taylor expansion, we can write this as:

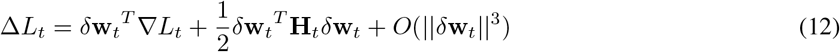

where **H**_*t*_ = ∇^2^*L*[**w**_*t*_] is the hessian of the task loss, which quantifies the effect of the local curvature of the loss landscape. For example, if *δ***w**_*t*_ projects onto directions of positive (upward) curvature, the second term will be positive and slow down learning (make Δ*L_t_* less negative).

Since Δ*L_t_* depends on the particular direction of the learning rule error 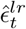, we look at the expectation over all the directions of 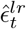. For the first term, using the fact that the online error 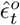 is perpendicular to the task gradient ∇*L_t_* and that on expectation the learning rule error is uncorrelated to the task gradient 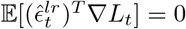 (see SI for full derivation), we get

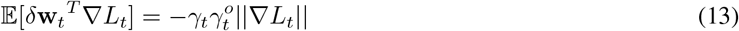

where 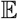 denotes the expectation over all the possible directions of 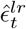. Equation (13), indicates that the change in task loss benefits from a steep slope, large ||∇*L_t_*||, and a small online learning error, large 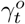.

For the second term, we can show that (see SI for full derivation)

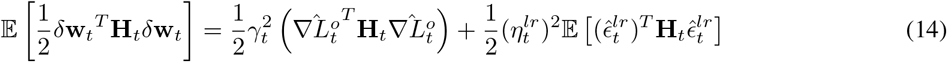

These terms capture the degree of curvature of the task loss landscape in the direction of 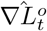 and 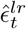 (first and second term respectively) at the point in weight space **w**_*t*_. This quantity depends on the eigenvalues of the hessian **H**_*t*_ and how strongly 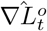 and 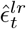 project onto the different eigenvectors of the Hessian. We can make this relationship concrete and approximate the dependency of the hessian projection terms on the eigenvalues of the Hessian^50^ (see SI).

The learning rule error 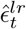 is uncorrelated with the task. Thus, in expectation it will project evenly onto the eigenvectors of the hessian (see SI). It follows that 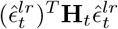 depends only on the average eigenvalue of the hessian (i.e. the average curvature)

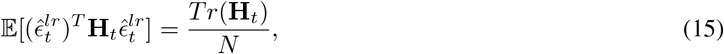

where *N* is the number of granule-cell-to-purkinje-cell weights and *Tr* denotes the trace of a matrix (Figure 3C left).

Unlike the learning rule error, the online gradient 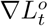 does not project evenly onto the eigenvectors of the hessian **H**_*t*_ (see Figure 3C right). In fact, as 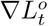 is a biased approximation of ∇*L_t_*, it tends to project more strongly onto the directions of large curvature of the hessian. We can bound the expected hessian projection above and below (see SI for full analysis):

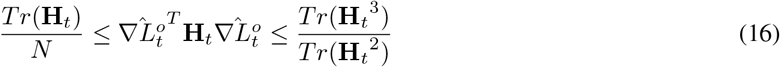

Where exactly 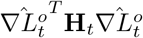 falls in the bounds depends on multiple factors: the weight state **w**_*t*_ and how close it is to a local minima, and on 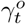. If the online gradient were uncorrelated with the task gradient, the hessian projection would be closer to the lower bound. As 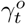 increases, then 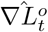 gets closer to 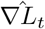 and 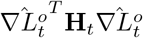 approaches the upper bound.

We can say that 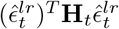 depends on the *average curvature* (Figure 3C left). As opposed to 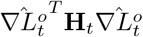 which depends on a *biased measure of curvature* of the task loss landscape (Figure 3C right). This biased measure depends on the spread of the eigenvalues of the hessian. Geometrically, this means that online learning suffers most when the loss landscape has highly anisotropic curvature.

### Effect of granule cell layer expansion on learning parameters

Our previous work showed that size expansions can mitigate the effect of loss landscape curvature on local learning rule performance.^51^ This suggests that problematic biases introduced by online learning can be similarly mitigated by adding granule cells in the model, as in Figure 2A.

This network expansion maintains the number of mossy fibre inputs *I* and increases the number of granule cells from *N* to *Ñ*. This increases the input expansion ratio from *N*/*I* to *Ñ*/*I*. Let 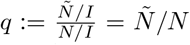 be the expansion ratio, defined as the ratio between the input expansion ratios of the two networks. Each new granule cell forms connections with *K* = 4 randomly chosen input mossy fibres. Hence the input weight matrix of the expanded network 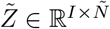 is composed of the initial weights matrix *Z* and added columns each with *K* non-zero elements.

We first analyse how online and task gradients are affected by this size expansion. These gradients depend both on the dynamics of the plant and the granule cell layer activity **h** from time 0 to *t* (see SI). The expansion of the granule cell layer does not affect plant dynamics. Hence, the change in the gradients from the expansion are determined by the change in **h**.

We can write the expected granule cell activity 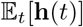 over a uniform distribution of the time points in the interval 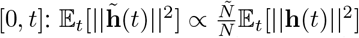. Consequently, the mean squared gradient will scale linearly with the number of granule cells. We may then write the normalised changes in gradient as:

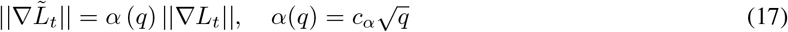

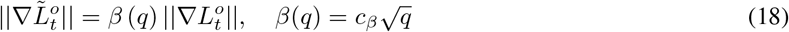

where 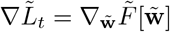 and 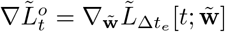 are the gradients in the expanded network and *α* and *β* are scalar functions that capture the scaling of the gradients as a function of the expansion ratio *q*. These calculations are backed by simulations (see SI Figure 1).

Next, we consider the scaling of the hessian projection terms with the network expansion. In the previous section, we stated that the projection of the online gradient onto the hessian can be bound by *Tr*(**H**_*t*_)/*N* and *Tr*(**H**_*t*_^3^)/*Tr*(**H**_*t*_^2^) as in equation (16) and the learning rule error projection only depends on *Tr*(**H**_*t*_)/*N* (see (15)). Thus, it suffices to evaluate how these three terms are modified with the network expansion.

Like the gradient, the hessian depends on plant dynamics and the granule cell representation (see SI). Without knowing the plant dynamics we cannot calculate directly the projections in the initial and expanded network or the hessian but we can consider how the quantities change relatively with the expansion (see SI). We find that the average eigenvalue stays constant 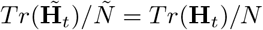 but the spread of the eigenvalues grows at most linearly 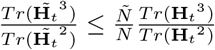.

Hence, the projection of the online gradient onto the hessian scales at most linearly with the network expansion.

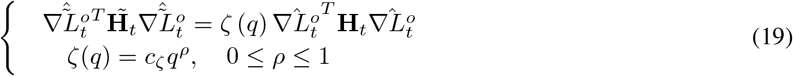

As mentioned in the previous section, 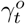 determines how close the projection is to the upper or lower bound. Hence, we expect *ρ* to increase with 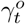. The more online error there is, the less accurate the online gradient approximation and the smaller *ρ* is (see SI).

The hessian projection of the learning rule error 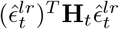 is equal to the average eigenvalue of the hessian (see (15)) which remains constant with the expansion

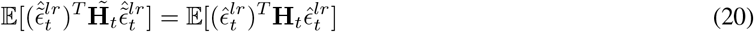

Taken together, we find that adding granule cells increases the slope of the task loss and the biased curvature, but preserves the average curvature. We next quantify the effects of these scalings on online learning performance.

### Trade-off between learning speed and steady state loss

Using the results from the previous section we can write the expected learning speed

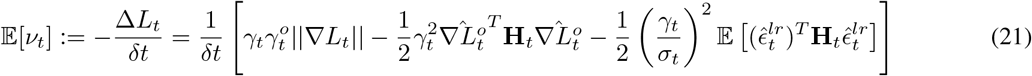

where 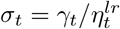 and *δt* is the time window between weight updates at time *t*.

Furthermore, we can define the expected local task difficulty as a proxy of steady state loss (see Methods and^51^):

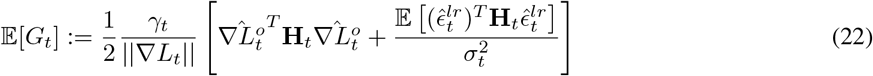

The local task difficulty quantifies the effect of the loss landscape on a change in weights *δ***w**_*t*_ near a local minimum.

A successful online learning system should simultaneously decrease rapidly the loss at initial parts of learning and achieve good performance at the end of learning. This is realized by a large learning speed 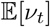 when the loss *L_t_* is large and a small local task difficulty 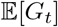 when *L_t_* is relatively small.

As illustrated by the motivating example, tuning the learning gain *γ* regulates the learning speed and the steady state loss. In fact, for fixed *σ_t_*, there is a learning gain that produces the optimal plasticity magnitude ||*δ***w**_*t*_|| in the direction 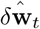 (see Methods)

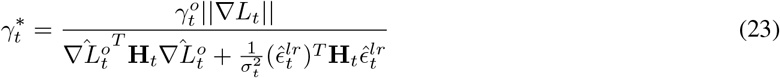

The optimal learning gain depends on both the quality of the learning rule and the local geometry of the loss landscape. In particular, 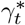 is proportional to the ratio between the slope ||∇*L_t_*|| and the hessian projections. At initial stages of learning, far from a local minimum, this ratio is large leading to large optimal gains. Near a local minimum, smaller gains are favoured as the slope is smaller and the upward curvature can lead to overshooting.

If at each learning step 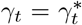, then both learning speed and steady state loss can be optimised. Biologically, it is unlikely that the system can perfectly compute 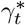 during learning because this would require access to the norm of the task gradient, the hessian projections and learning rule error throughout learning. We therefore set *γ* to be constant during learning and consider the effect of circuit architecture on overall learning performance for a range of fixed *γ* that avoids instabilities at one extreme and arbitrarily slow learning at the other. We will pay particular attention to the best performing value of *γ* to assess the impact of size expansion because this corresponds to the case in which the biology is doing the best it can do with synaptic plasticity.

If *γ* is very large relative to 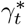, learning is unstable —the task loss does not decrease or reach a steady state value. On the other hand, a small *γ* leads to a slow learning speed but better steady state performance. In the range of gains that guarantee stability, we find there is a trade-off between gains that favour learning speed and gains that favour steady state loss (Figure 4A).

**Figure 4:**
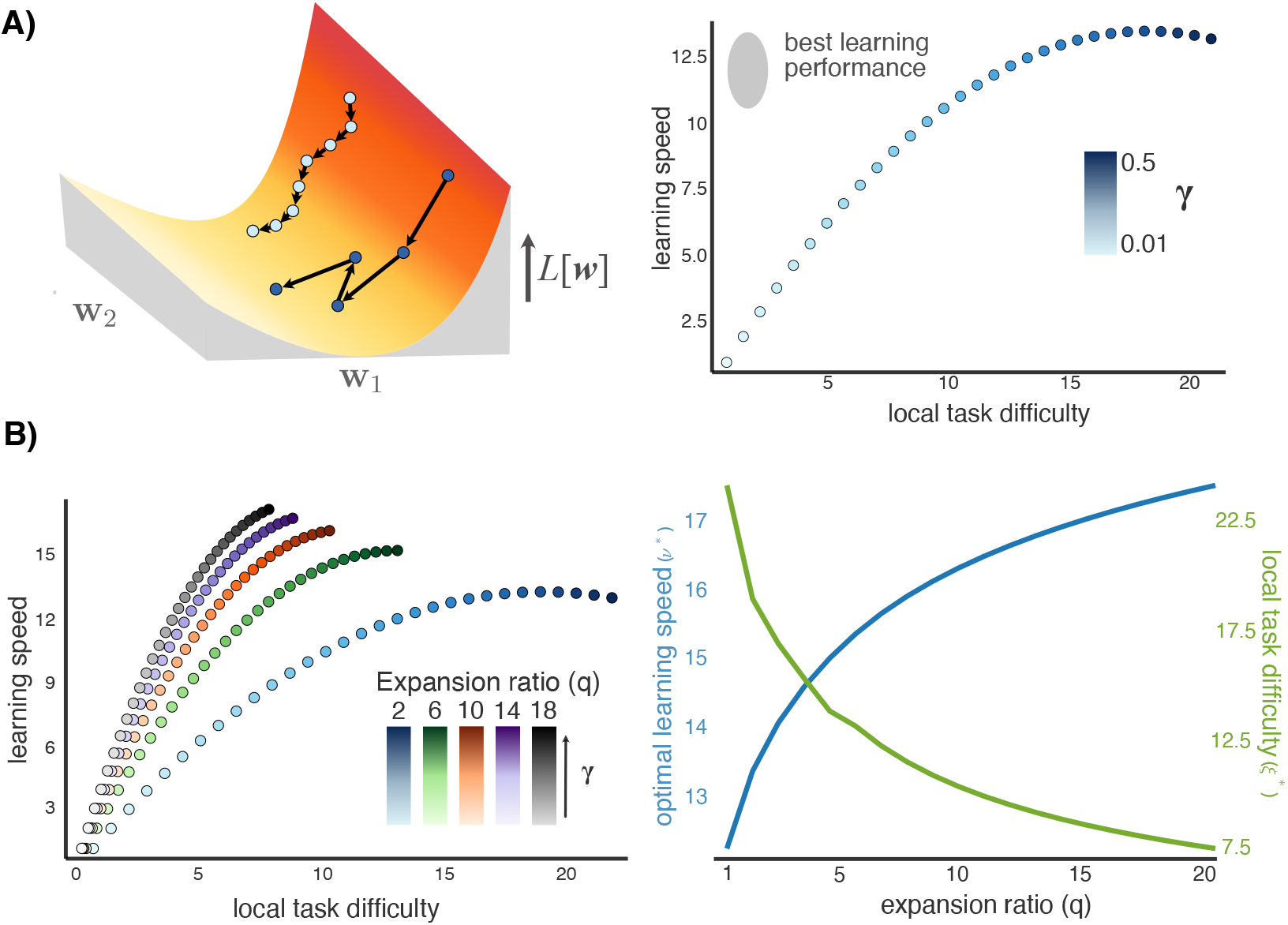
The granule cell layer expansion navigates the trade-off between learning speed and steady state loss. **A)** The strength of *γ* represents the sensitivity of the learning rule to new information. This quantity modulates a trade-off between learning speed and steady state loss. On the left, visualisation of a weights trajectory during learning for a small *γ* (light blue) and large *γ* (dark blue). A small *γ* descends the loss landscape slowly (small learning speed) but can get closer to a local minimum (small steady state loss). A large *γ* leads to large steps that descend the loss landscape fast but oscillate around the local minimum. On the right, steady state loss vs learning speed for different values of *γ*. **B)** Adding granule cells mitigates the trade-off between learning speed and steady state loss. On the left, steady state loss vs learning speed for different network expansion values *q*. The different colors correspond to different *q* and the variation in shade codes for the change in *γ*. As *q* increases, the system can achieve better learning speed for the same steady state performance. On the right, the steady state loss (in green) for the maximal learning speed (in blue) decreases with *q*, hence better learning speed and steady state loss can be achieved simultaneously by increasing the input expansion.

### Granule cell expansion navigates the trade-off between learning speed and steady state loss

Tuning the gain *γ* for optimal learning introduces a trade-off between learning speed and steady state performance. We now show that by modifying the loss landscape, the granule cell layer expansion mitigates this trade-off, improving performance in both measures regardless of where the biology has chosen an acceptable balance.

First, we look at how expanding the granule cell layer modifies the learning speed in (21). Using (17), (19), and (20), we find that in the expanded network

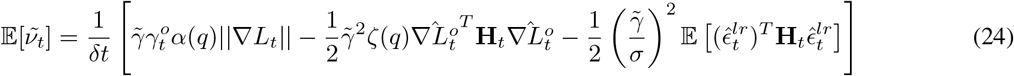

As 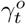 and *σ* do not depend on the network architecture, only on the quality of the learning rule and the limitations form online learning, we have 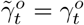 and 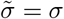. However, networks with different expansion ratios could have different learning gains 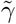. We show that for any *γ* in the initial network, we can find a 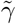 in the expanded network that improves learning speed 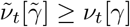 (see Methods).

Second, we can express the local task difficulty as defined in (22) in the expanded network

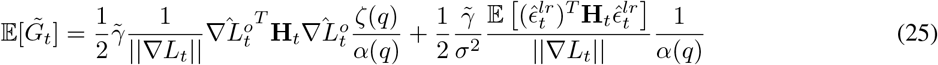

As the slope increases but the average curvature is constant, the second term decreases with the expansion. We can show that there is a 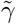 in the expanded network that reduces the overall local task difficulty 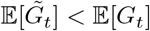 (see Methods).

Combining the two 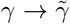 mappings that improve learning speed and local task difficulty, we get a formula for the mapping that improves both simultaneously

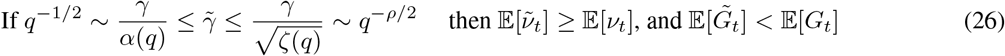

Increasing the input expansion and reducing *γ* can take advantage of the change in loss landscape to to improve overall learning performance. In theory, the learning performance can keep improving with an increasing expansion ratio, as long as the gain decreases in parallel. However, in practice, very small gains cannot be implemented biologically as a small change in synaptic plasticity could be out-weighted by synaptic noise.^51^ This provides a lower bound on *γ* and the corresponding maximum input expansion *q* for a learning problem.

The mapping of 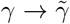 is determined by the scaling functions of the gradient *α*(*q*) and of the hessian projection of the online gradient *ζ*(*q*). As mentioned in previous sections, the scaling of the hessian projection depends on 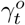 —the correlation between the online gradient and the task gradient. In particular, the scaling factor *ρ* decreases with 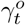 (see SI). It follows that learning problems with high online learning error (small 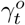) can benefit from larger input expansions (see supplemental Figure 2).

We have shown that the granule cell layer expansion modifies the geometry of the loss landscape in a way that benefits online learning. For any expansion ratio, there is a trade-off between learning speed and steady state loss modulated by *γ*. However, increasing the expansion ratio by adding granule cells relaxes the trade-off allowing the networks with larger input expansions to achieve better learning speed and steady state loss simultaneously (Figure 4B).

### Granule cell activity density affects learning performance

Having established that online learning benefits from an expansion layer, we hypothesized that the benefit of this expansion should depend on the fraction of cells that are active during learning. The benefit of a size expansions requires the sensitivity of task error to be distributed across population activity. Cells that are silent do not carry such information and therefore reduce the effective size of the expansion.

Granule cell activity in the expansion layer depends on the activation function *ϕ* in eq (2). In the previous sections, we assumed smooth activation functions, but this means there is no definite threshold at which a unit transitions from being inactive to active. We therefore replaced the smooth (tanh) activation function with a rectified linear unit (reLu):

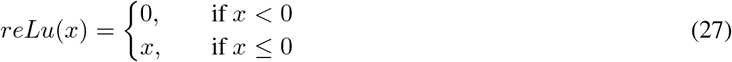

as depicted in Figure 5A. Unlike tanh, *reLu* is identically zero beneath threshold and has graded activity above this threshold. This lets us define population activity density, *f*, as the average fraction of active granule cells throughout the learning task. This measure is called the ‘coding level’ in other work.^3,52^

**Figure 5:**
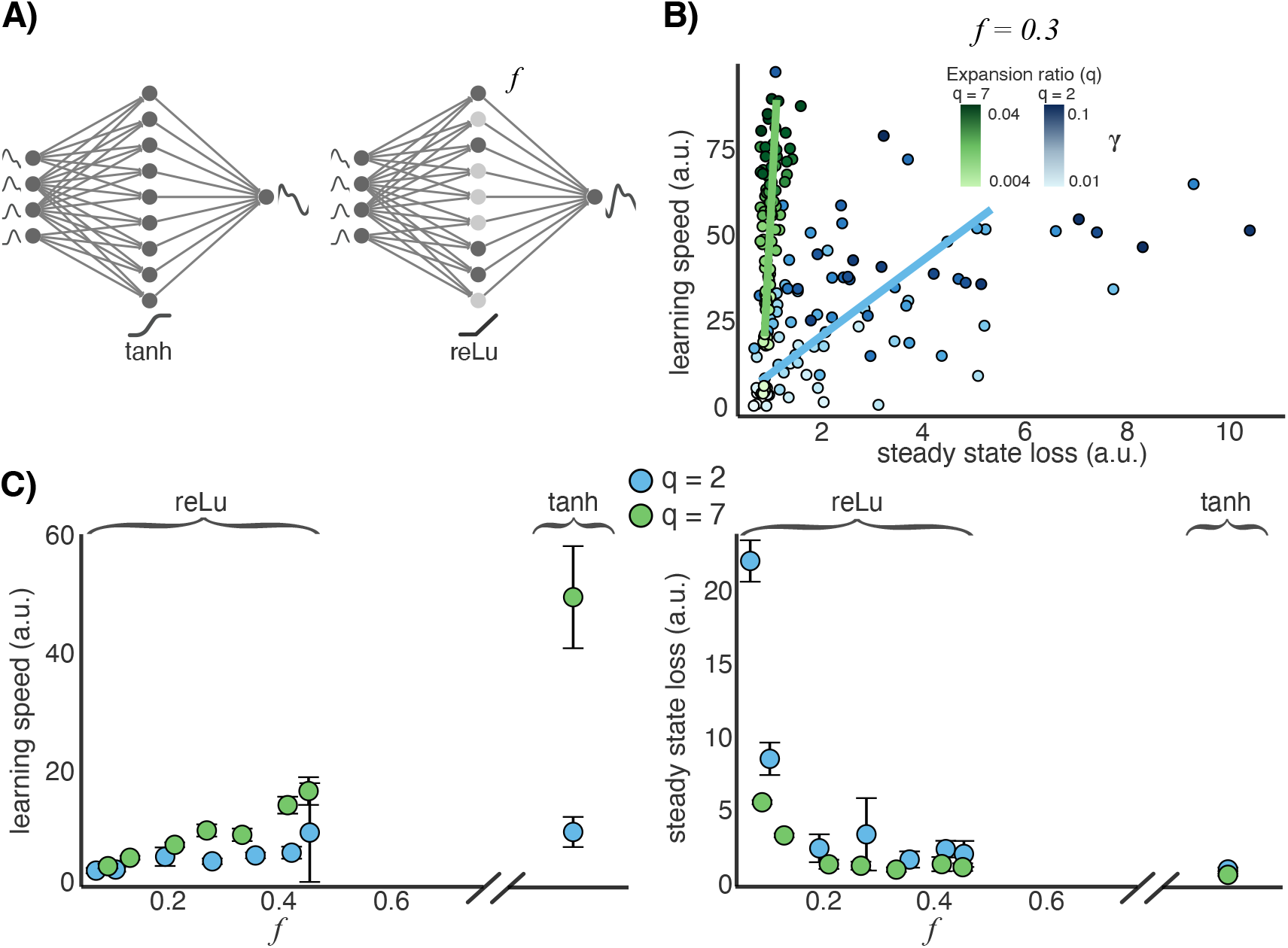
The effect of granule cell layer activity density on learning performance. **A)** We test two different activation functions for the granule cells: tanh (on the left) and reLu (on the right). With *reLu* some granule cells are not active (light colored cells). *f* represents the average fraction of active granule cells during the whole trajectory. **B)** The effect of input expansion for sparse activity density *f* = 0.3. We plot the trade-off between learning speed and steady state loss for two different expansion ratios: *q* = 1 in blue, *q* = 7 in green. The shade of the color denotes the value of *γ*. **C)** We plot the learning speed and steady state loss for networks with reLu activation function and varying *f* and networks with tanh activation function. In each case we test two different expansion ratios (*q* = 2 in blue and *q* = 7 in green). On the left, the learning speed increases with *f*. On the right, the steady state loss decreases with *f* the effect saturates for larger *f*. In both cases, the tanh activation function, which has fully dense coding, achieves optimal learning performance.

We systematically adjusted *f* by changing the distribution from which we draw the biases of the granule cells *b*_1_, *b*_2_,…, *b_N_* (see Methods). We first verified that the results of previous sections apply for this non-differentiable activation function. Both the trade-off between learning speed and steady state loss, and the benefit of a size expansion hold for any fixed *f* (Figure 5B).

We then assessed whether modifying *f* itself affects learning performance. We found that both learning speed and steady state loss benefit from denser granule cell layer activity independently of the input expansion (Figure 5B). Intuitively, the optimal learning performance is comparable to that obtained with a smooth activation function. This is expected because in the smooth case there is no sharp threshold that can silence cells and effectively reduce the population size. Furthermore, the activity density *f* affects learning speed and steady state loss differently. Learning speed increases linearly with *f*, while steady state loss benefits from even a slight increase in the density at low values of *f*. This benefit plateaus at a fraction, *f* = 0.2, above which the steady state performance is relatively insensitive to *f*.

We saw that changing the input expansion in the cerebellar network does not affect the amount of online learning error and learning rule error in the learning rule, but modifies the loss landscape in a way that benefits online learning performance. Unlike the input expansion, the granule cell layer activity density *f* modifies both the loss landscape and the relative amount of online learning error and learning rule error. The slope of the loss landscape is proportional to 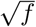, and thus increases with the density of the granule cell layer activity, accounting for the improvement of learning speed with *f*. Furthermore, sparser activity leads to larger online learning error, consistent with the hypothesis that reducing the number of active cells results in a smaller usable fraction and a smaller effective size expansion.

## Discussion

Half a century ago, Marr and Albus proposed a computational model of cerebellar function in which the granule cell layer separates input activity patterns.^1,2^ Subsequent work broadened the study of expansion recoding for pattern separation and explored its implementation in a variety of complex models.^3,4,7–9,53–57^

The classical theory of expansion recoding in the cerebellum relies on sparse activity in the granule cell layer. However, some studies of granule cell activity suggested that they might not always act as sparse coding pattern discriminators.^58^ Furthermore, multiple recent experimental studies found that the granule cell population activity is much denser than previously believed.^12–16^ These findings rekindled theoretical work on the granule cell representation for pattern separation.^3,4,7^ The work found that population sparsity might not be necessary for efficient pattern separation. In fact, decorrelation at the granule cell layer is the main determinant of pattern separation performance.

Our analysis dropped the assumption of pattern separation altogether and established a role for size expansions in reducing biases caused by online learning. Crucially, we found that denser granule cell activity improves both learning speed and steady state loss but has a greater benefit for fast learning. Furthermore, the effect of the input expansion on the trade-off between learning speed and steady state loss is conserved for different coding levels. Recent unpublished computational work corroborates our study, showing that for continuous input-output mappings, denser representations achieve better steady state loss when trained with gradient descent^52^ and that dense activity favours time-series learning.^59^

Though we aimed at generality, it is important to acknowledge the diversity of experimental findings as well as some of the specific choices we made in this study. Granule cell firing rates are found to vary across cerebellar regions.^60,61^ In line with the distinction between static pattern separation and online learning of dynamics, these differences might reflect involvement in different type of tasks requiring different coding levels. Our analysis of how loss functions are affected by by size expansions implies that cerebellar architecture is beneficial for learning with limited information, regardless of coding level.

While is widely accepted that the cerebellum acquires internal models for motor control and other dynamic cognitive tasks,^62–67^ the exact organization of internal models is still under debate.^11,32^ We chose a model in which the cerebellum learns an inverse model that assists in realising a desired reference trajectory. It is also believed that the cerebellum can learn a forward model that maps an efference copy of the motor command to the estimate of the plant state.^27,31,32,68–70^ In fact, both forward models and inverse models can be viewed as inter-related. Acquisition of a motor act requires inverse models, but the acquisition of the models itself requires forward models. We expect results to apply for a forward model because the difficulties posed by online learning are analogous. Although the signals and loss functions differ, the decomposition of the change in weights (10) and the effect on the loss landscape from the expansion still applies.

There are other limitations to our work concerning the level of biological detail. We used a firing rate model and a simple musculoskeletal plant. These choices made analysis more tractable and, we hope, our results more general. Other simulation based studies have used spiking models and complex musculoskeletal systems.^71–74^ These studies show that the basic architecture of the model we considered learns in a similar way despite such complexities. We modelled a cerebellar-like network as a three-layer feedforward network. This description is a simplification which omits interneurons and deep cerebellar nuclei. Golgi cells are excited by mossy fibres and parallel fibres and inhibit granule cells.^75,76^ Molecular layer interneurons provide inhibitory influence to purkinje cells’ dendrites which can have a strong effect on the cerebellar cortex computations.^77,78^ Interestingly, inhibition in the input layer of the cerebellar cortex can decorrelate and sparsen granule cell activity.^3,8,79^

Our analysis shows how online cerebellar learning imposes constraints on the quality of feedback information. Anatomically, such information is believed to reside in climbing fibre signals, but the physiological events that convert putative error signals into synaptic changes are complex. Ito proposed that parallel-fibre-to-Purkinje-cell synapses undergo long-term depression (LTD) when climbing fibres and parallel fibres are activated.^80^ Long-term potentiation (LTP) was later discovered to lead synaptic plasticity in the absence of climbing fibre inputs.^81^ Since then, multiple forms of plasticity have been found at almost every synapse in the cerebellum circuit.^82–86^ We remained agnostic of the exact learning rule while acknowledging biological limits: locality in both space and time, finite precision, delays, inability to optimise learning rates online. It would be particularly exciting to find empirical evidence that these general assumptions are strongly violated.

The diversity of physiological and anatomical details reinforces the view that one should consider the collective effect of plasticity mechanisms^87^ and how these interact with circuit architecture^47^ to understand overall learning performance. In putting forward the Levels of Analysis paradigm, Marr stressed the need to account for hard *implementational* constraints coming from cellular physiology when analysing an *algorithmic* means of solving an overall *computational* function.

In the present study, we could not explain the role of cerebellar connectivity without simultaneously considering the computational task of dynamic learning and the low-level characteristics of synaptic plasticity and population sparseness. Our reading of Marr is thus informed by one of the more general lessons we learned from conducting this study: levels of analysis cannot be applied in isolation while remaining biologically relevant.

## Methods

Full details of all simulations are provided in the *Supporting Information*. In this section we provide an overview. All the simulations were done on Julia.

### Motor control system

We model the motor control system in Figure 1A.

**The feedback controller** is modelled as a PID controller with gains *K_p_* = *K_d_* = *K_i_* = 0.1. The output is given by 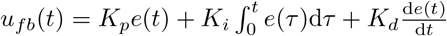, where *e*(*t*) = *r*(*t*) − *y*(*t*) is the trajectory error.

**The musculoskeletal system** is modelled as a single input single output stable linear plant with the space state model

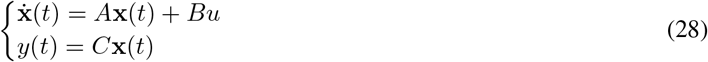

where

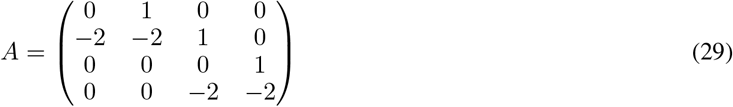

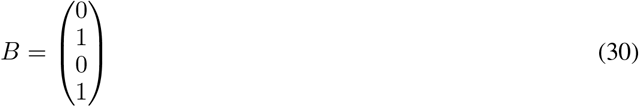

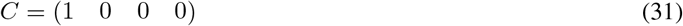

For simulations we chose the initial conditions to be all zeros.

**The reference trajectory** is the *r*(*t*), *t* ∈ [0, *T*] that the system needs to learn to track. In any system there are signals that the plant is not able to produce because of the limitations of its dynamics. Biologically, this makes sense as there are some movements that are too fast or to big for the musculoskeletal system to produce. We can express this limitation by a cut-off frequency *f_c_*. We assume that any signal that has frequencies below *f_c_*, can be actuated by the plant. We chose *f_c_* = 0.1*Hz*. We want an unbiased reference signal that can be learned by the system.

We define the reference signal from a Fourier series as follows

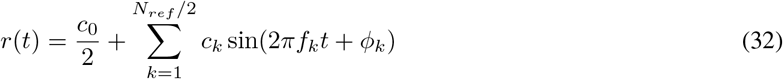

where *N_ref_* is the number of different sinusoidals used to generate the signal. We choose by default *N_ref_* = 200. The harmonic frequencies are

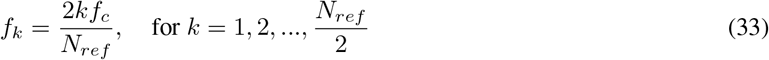

This satisfies that the highest harmonic frequency matches the cut-off frequency *f_c_*. The phase shifts *ϕ_k_* are drawn from a random uniform distribution over the interval [0, 2*π*). The constant coefficients *c_k_* are selected according to the desired power spectrum. We want to approximate a filtered Ornstein-Uhlenbeck (O-U) process. The power spectrum of an O-U process is a Lorentzian. Hence we draw the *c_k_* coefficients from a Lorentzian

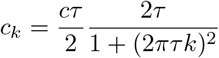

where *τ* is the inverse of the width and *c* controls the magnitude of *c_k_*.

We want |*r*(*t*)| < 1 to make sure that the plant can realize the reference trajectory. Yet the steady state variance of an O-U process with coefficients *c* and *τ* is 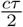. We chose 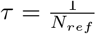 and 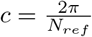.

**The inputs to the cerebellar-like network m**(*t*) are defined as in eq (1), where Δ*t_I_* = 0.2/*f_c_* and *δt_I_* = Δ*t_I_*/(*I* − 1) where *I* is the number of inputs to the cerebellar like network. Learning is relatively sensitive to the choice of Δ*t_I_*; if it is too small, there is no enough lookahead for the cerebellar like network to learn the inverse model. If *δt_I_* is larger than one quarter of the period of the largest frequency sinusoidal, the inputs **m** do not reflect the dynamics of the reference signal hence learning night not happen. We define the length of the lookahead window Δ*t_I_* with respect to the cut-off frequency *ω_c_* to make sure that the lookahead window is large enough to include some variation in the reference trajectory. Our results are robust to the choice of these parameters as long as the lookahead window includes a non-constant portion of the reference trajectory [*r*(*t*), *r*(*t* + Δ*t_I_*)] but no more than quarter of the period of the largest frequency sinusoidal.

**Cerebellar-like network** For simulations, we initialise the network architecture to have *I* = 10 inputs, a single output and the number of granule cells *N* varies. Each granule cell passes inputs through an activation function *ϕ*. For all simulations, except in the last results section, we use the hyperbolic tangent non-linearity tanh : ℝ → ℝ of the form 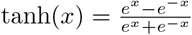. The Purkinje cells have a linear output.

Each granule cell receives *K* = 4 inputs from randomly chosen mossy fibres. The input weights are drawn from standard-normal distribution with mean 0 and standard deviation 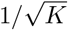. Hence, the input weight matrix *Z* has *N* columns with each column having *K* non-zero elements. The input weights are constant throughout the trajectory.

The biases *b_i_* are drawn from a random uniform distribution 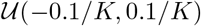, except for the last results section where we draw from different uniform distributions to control the coding level. In the last results section, we select different values of *θ* = −1, −0.5, −0.1, 0, 0.1, 0.5, 1 and draw the biases from the distributions 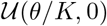, if *θ* < 0 and 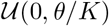, if *θ* > 0.

The granule cell layer to purkinje cell layer is fully connected. Each output weights is drawn from a standard normal distribution with mean zero and standard deviation 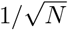. The output weights are plastic and are updated according to a learning rule.

**Learning rule** The goal of this paper is not to determine learning rules in the cerebellar cortex for motor control. However, we test different learning rules to to determine the validity of our model. The weights are updated throughout the trajectory every *δt_e_* = 1, we set this interval to be uniform for convenience. We use an adapted version of least mean squares (LMS).^48^ The weight update at each update time is given by (5). The vector of learning rule error 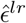, each element is drawn independently from a random normal distribution 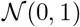.

The classic LMS rule is given by *δ***w**_*t*_ = −*γ***h**(*t*)(*r*(*t*) − *y*(*t*)). The main differences between our learning rule and classic LMS are

- In classic LMS the weight update is continuous, in our case, the weights are updated at discrete points in time during the trajectory.
- The climbing fibre activity encodes the trajectory error over the time window [*t* − Δ*t_e_* − Δ*t_r_, t* − Δ*t_r_*], instead of the instantaneous error. Δ*t_r_* denotes the delay between the trajectory is executed and the climbing fibre signal arrives to the cerebellar network.
- We introduce a possible delay between the granule cell activity and the climbing fibre activity leading to synaptic plasticity. This delay is represented by Δ*t_h_* (see^45^).
- We include learning noise 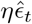 arising from imperfections in the synaptic plasticity mechanisms or in the information on the error *e*(*t*).

Unless otherwise specified we set Δ*t_e_* = 1.0, Δ*t_r_* = 0.5 and Δ*t_h_* = 0.1.

### Network expansion

The network expansion maintains the number of mossy fibre inputs *I* and increases the number of granule cells from *N* to *Ñ*. This increases the input expansion ratio from *N*/*I* to *Ñ*/*I*. Let *q* ≡ *Ñ*/*N* be the expansion ratio, defined as the ratio between the input expansion ratios of the two networks. Each new granule cell forms input connections with *K* = 4 randomly chosen input mossy fibres. Hence the input weight matrix of the expanded network 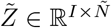 is composed of the initial weights matrix *Z* and added columns each with *K* non-zero elements.

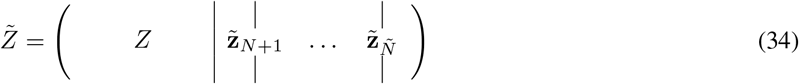

Each added granule cell forms a connection with the output purkinje cell. We assume the new connections initially have weight zero so the initially do not contribute to the input-output properties of the network. The output weights of the expanded network are

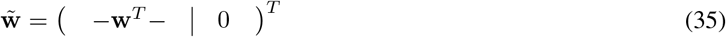

This expansion preserves the input-output properties of the cerebellar-like network before learning. Indeed the task loss of the original and expanded network are equal 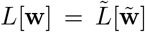 but the granule cell layer activity **h** changes 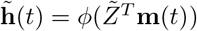.

### Details for simulations in motivating example

We simulate the system as described in the results for a trajectory time *T* = 1000. At each time of weight update *t*, the task loss *L*[**w**_*t*_] is computed. To do so, a system identical to the one being trained, except with fixed output weights **w** (no weight update) is defined. We compute the task loss as defined in eq (6) in this static system. Indeed, the task loss reflects the performance of the output weights **w** on the task, over the whole trajectory. The task gradient *L_t_* is also computed in the static system.

For Figure 2, we fit the task loss *L*[**w**_*t*_] with an exponentially decaying function 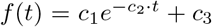. We take the **learning speed** *ν* = *c*_2_.

To compute the **steady state loss** we pre-train the network with gradient descent to a very low task loss (we set *T* = 7000 and *γ* = 0.1 for pre-training). The weights after pre-training are close to the optimal weights. We use these weights as initial weights and train with the LMS-like learning rule. During this training, the task loss initially increases—as the initial weights are very close to the local minimum—and then settles around a steady state value. We set the steady state loss to the mean of the task loss over the last 100 steps. The pre-training is important to recover the optimal steady state value without having to train the network with the LMS-like learning rule for a very long trajectory time. In the absence of pre-training the systems, especially trained with small *γ* might not reach steady state.

We compute the learning speed and steady state loss values for a range of number of granule cells *N* = 10,…, 90 and a range of *γ*. We perform the network expansion as described in the results. For each network expansion explore a different range of *γ* indeed, larger networks in general require smaller *γ* for learning. For Figure 2, we have 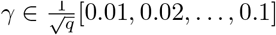. We normalise the values of the learning speed and steady state loss by the values for the smallest *q* and *γ* to emphasize the relative changes.

For Figure 2, we performed 25 simulations with different reference trajectories and different weight initialisations for the cerebellar like network. We selected the simulations with similar initial task loss (loss before learning). In particular, we discard simulations with initial loss below the average initial loss minus 1/2 standard deviation. This filters out initialisations with very low initial loss that might not converge when trained with the upper range of *γ*. After filtering out we are left with 17 simulations.

For Figures 2C and D, for each simulation, and for each *q*, we compute the optimal learning speed over all *γ* and the steady state loss value for the same value of *γ*. Each point represents the mean over all simulations and the error bars represent the standard error.

Figure 2B represents the trade-off between steady state loss and learning speed over *γ* for a single simulation. For one reference trajectory, we train for all *N* and *γ* starting at the same weight initialisation 20 different times. The learning rule error 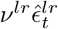 in the weight update (5) introduces a slight variability each training simulation. We plot the learning speed vs steady state loss for *q* = 1, 7, for all *γ* and for all training simulations. We plot the trend line for each *q*.

For more details on these simulations see SI.

### Measuring learning parameters

For Figure 4, we compute expected learning speed 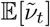 and local task difficulty 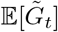 as a function of expansion ratio *q* (see eq (24) and (25)). We first compute the value of the learning parameters and scaling functions found in the equations (24) and (25).

We train the motor control system as described in the results with a weight update given by (8) with Δ*t_e_* = 1.0 and Δ*t_r_* = 0.5. The online gradient 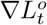 is computed using local sensitivity analysis. Note that the gradient has to be taken across the system, including the plant, proportional-integral-derivative (PID) controller and cerebellar-like network. Julia is able to compute the gradient across the whole system. Each element of learning rule error 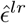 is drawn independently from a random normal distribution 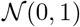.

At each weight update time, we compute the task gradient 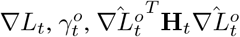 and 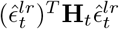.

We generate expanded networks, with expansion ratios *q* = 1, 2,…, 11. In each expanded network, we compute the learning parameters as described above. We approximate the scaling functions *α*(*q*), *β*(*q*) and *ζ*(*q*) fitting the variables with respect to the expansion ratio. We use a fit of the form 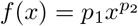. In particular, we can get an estimate of *ρ*, the scaling of the biased curvature.

For Figure 4, we use the scaling functions and the learning parameters found from the simulations to compute the expected learning speed 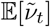 and local task difficulty 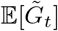 as a function of expansion ratio *q* (see eq (24) and (25)). We use these equations to compute the learning speed and local task difficulty as we vary the expansion ratio *q* and the *γ*. We test *q* = 1, … 20 and 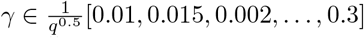. We normalise the values of *ν* and *G* for each *q* and *γ* by the value for *q* = 1 and smallest *γ*.

### Granule cell layer activity density simulations

For the simulations on the granule cell layer activity density, the granule cells have reLu activation function. We control the coding level of a system through the distribution from which we draw the biases. The fraction of non-zero granule cells varies along the trajectory. We compute the coding level as the mean fraction over the whole trajectory.

For Figure 5 we run simulations as described above. For Figure 5A we simulated for a single reference trajectory and different bias distributions and train each system 10 different times. The scatter plot shows the results for each simulation.

For Figure 5B, we average the learning speed and the steady state loss for 4 simulations with different reference trajectories and weight initialisations, training each for different distributions for the biases. We train each system 5 times and average over the instances.

### Steady state loss analysis

Consider the expected change in task loss 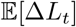, at some time *t* given a change in weights *δ***w**_*t*_. We can rewrite the expression in eq (12) as follows

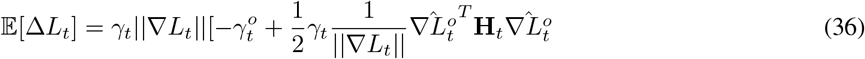

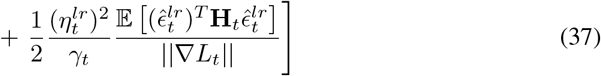

Learning requires a negative change in task loss, and it stops when

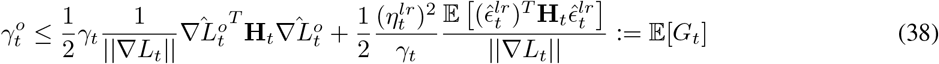

We refer to the right hand side of this equation as the local task difficulty based on the definition by Raman et al.^51^ This inequality is bound to be satisfied as when the task loss approaches a local minimum ||∇*L_t_*|| goes to zero, **H**_*t*_ becomes positive definite hence the hessian projections 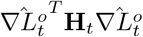 and 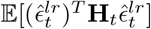 are positive.

### Optimal plasticity magnitude

Given a direction of weight plasticity 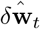, we can find the optimal plasticity magnitude ||*δ***w**_*t*_||* that minimises the change in task loss Δ*L_t_* given in eq (12) (see Methods and^50^)

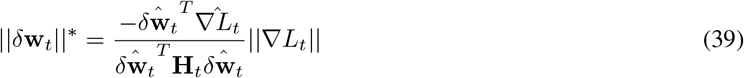

(as long as 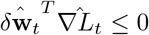, and 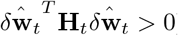). The optimal plasticity magnitude depends linearly on the magnitude of the gradient (i.e. the slope of the task loss surface) and is decreased by the projection of 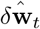 on the hessian of the task loss. In general, far from a local minimum (i.e when the gradient is large with respect to the curvature) the optimal plasticity magnitude is large. During learning, the gradient norm decreases with respect to the curvature hence the optimal plasticity magnitude decreases. Close to a local minimum, a large change in weights can lead to overshooting preventing a decrease in task loss (see Figure 4A).

With fixed 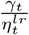 we can find the optimal learning gain 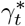 such that 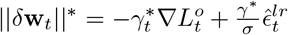. Using this with eq (39) we get

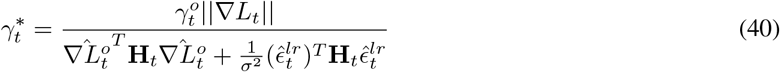

If *σ* is large (i.e. small learning rule error), then the second term in the square root is small and we get 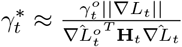.

The learning rule error effectively decreases 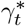.

At initial stages of learning, ||∇*L_t_*|| is relatively strong compared to the hessian projections, and decreases with learning. Hence *γ*\* is large initially and decreases as the task loss decreases.

Close to a local minimum, as the task gradient converges to zero, the *γ*\* also converges to zero. Indeed, if the weights were at the exact local minimum the optimum change in weights would be zero. In practice, however, it is unexpected for a learning rule with errors to converge perfectly to the local minimum. Hence, *γ*\* in practice will be non-zero.

In general, the *γ* that optimises learning speed 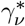 is larger than the one that optimises steady state error 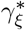.

For a fixed *γ* over learning, there are two regimes

- 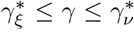: there is a trade-off between learning speed and steady state loss. Increasing *γ* makes steady state loss worse but learning speed better.
- 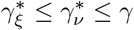: both learning speed and steady stater error get worse, learning can reach instability

As in the second case there might not be any learning because of instabilities, the first case is the relevant one for this analysis.

### Learning speed scaling

We can show that there is a mapping from 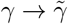 that guarantees an improvement in the learning speed. The first term in 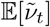 (24) increases if and only if

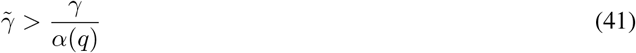

The second term decreases if and only if

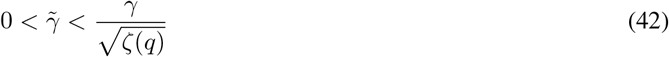

And the third term decreases if and only if

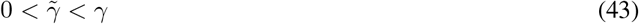

From the scaling of the learning parameters we have *α*(*q*) ~ *q*^1/2^ and 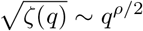 with 0 ≤ *ρ* ≤ 1. Yet *q* ≥ 1 hence *q*^−1/2^ ≤ *q*^−*ρ*/2^ ≤ 0. Thus,

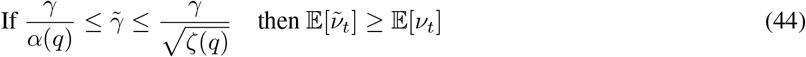

with the equality happening if *ρ* = 1.

### Steady state loss scaling

The first term in the local task difficulty (25) decreases with the expansion if and only if

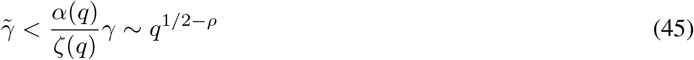

and the second term decreases with the expansion if and only if

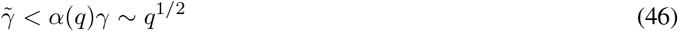

Putting these two equations together we get

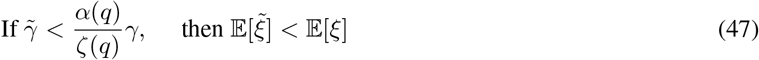

We can find *γ* in the expanded network such that the steady state loss is reduced by the network expansion.

## Acknowledgments

This work was supported by ERC grant No. 716643 FLEXNEURO.

## Author contributions

Conceptualization, A.P.R., D.V.R., and T.O.; Methodology, A.P.R., D.V.R., and T.O.; Investigation, A.P.R.; Writing— Original Draft, A.P.R; Writing—Review & Editing, A.P.R., D.V.R., and T.O.

## Declaration of interest

The authors declare no competing interests.

## Supplemental Information

### Expected change in task loss

Consider the system at some time *t* during the trajectory. The cerebellar weights have a value **w**_*t*_ and the change in weights from the learning rule is *δ***w**_*t*_. The change in task loss Δ*L_t_* from the weight update *δ***w**_*t*_ is

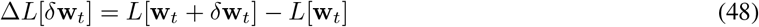

If the system is learning, i.e. the weight update is beneficial, then Δ*L*[*δ***w**_*t*_] is negative. A large change in loss at the beginning of learning leads to a large learning speed. The steady state is determined by the loss when the change in task loss becomes zero or negative.

We can use the Taylor expansion on the task loss *L*[**w**_*t*_ + *δ***w**_*t*_] to rewrite the change in task loss

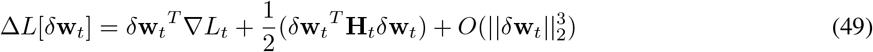

where ·^*T*^ corresponds to the transpose of a vector and we have used the shortened notation for the task loss gradient *L_t_* ≡ ∇_**w**_*L*[**w**(*t*)], and hessian of the task loss 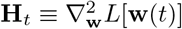. The first term is the contribution from the gradient. It quantifies the correlation between weight update from the learning rule and the perfect gradient descent. The second term quantifies the contribution of the curvature of the task loss surface. In particular, it is proportional to the degree of curvature in the direction of the weight change **w**. The smaller this term is, the larger loss change and better learning performance. Note that the last term 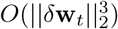 might not be small, but if we consider small time steps between weight updates then this term will be small compared to the first two. Hence from now on we use the second order approximation of Δ*L_t_*.

Using the weight update eq (8) in the second order approximation of Δ*L_t_* we get

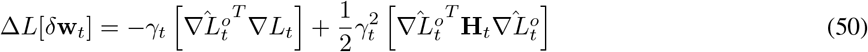

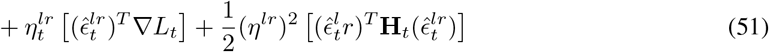

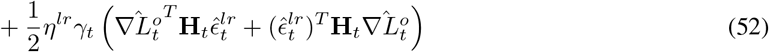

where 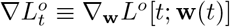 is the shortened notation for the online gradient. The first two terms are the contributions from the task relevant plasticity (i.e. the change in loss in the online gradient direction). The next two terms are the contribution from the learning rule error.

In expectation (over the learning rule error directions), 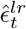 is uncorrelated with the derivatives of the task loss ∇*L_t_* and **H**_*t*_, otherwise it would be a task-relevant term. Hence, on expectation

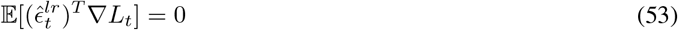

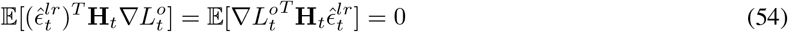

Hence on expectation the change in loss is as follows

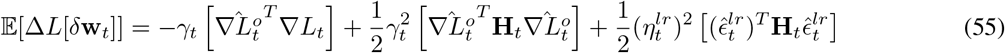

### Hessian projections

Consider the hessian terms 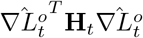 and 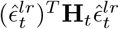. These terms capture the degree of curvature of the task loss *L* in the direction of 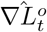 and 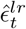 at the point in weight space **w**_*t*_. This depends on the eigenvalues of the Hessian **H**_*t*_ and how strongly the online gradient 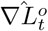 and the direction of learning rule error 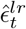 project onto the different eigenvectors of the Hessian.

We can make this relationship concrete and approximate the dependency of the hessian projection terms on the eigenvalues of the Hessian.^50^

First, consider a random vector *E* uncorrelated with the task loss. Let 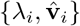, *i* = 1,…, *N* be the eigenvalue-eigenvector pairs of the hessian **H**_*t*_. By symmetry of **H**_*t*_, we can construct an orthonormal coordinate system from the eigenvectors of the hessian 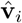. On expectation, it projects evenly onto all directions of the eigenvectors 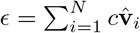. It follows that the projection onto the hessian

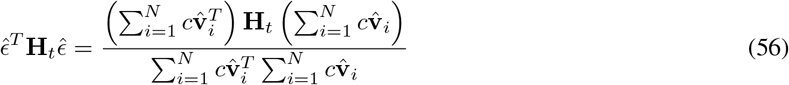

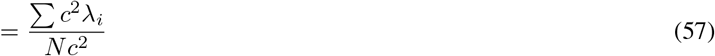

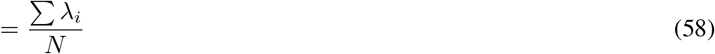

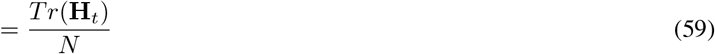

where *Tr*() denotes the trace of a matrix. In the last line we used an algebraic property of the relationship of the eigenvalues of a matrix and its trace. Hence, a random vector, uncorrelated with the task loss, projects evenly onto all the eigenvectors of the hessian. Its projection is onto the average curvature 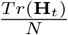.

The learning rule error 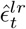 is uncorrelated with the task. It follows that 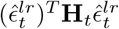 depends only on the average eigenvalue of the hessian (i.e. the average curvature)

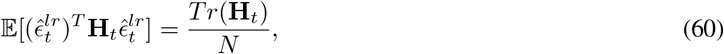

Before looking at the projection of the online loss gradient 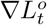, we consider the projection of the task loss gradient ∇*L_t_*. Unlike the learning rule error, the task loss gradient ∇*L_t_* does not project evenly onto the eigenvectors of the hessian **H**_*t*_. Consider the loss landscape near a local minimum **w**\* The loss landscape can be approximated by a quadratic function in the weights **w** and the hessian **H**_*t*_ is constant and positive definite. In this case, ∇*L_t_* = **H**_*t*_(**w**_*t*_ − **w**\*). The vector **w**_*t*_ − **w**\* can be decomposed in the directions of the eigenvectors of the Hessian 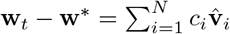. Where 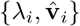, *i* = 1,…, *N* are the eigenvalue-eigenvector pairs of the hessian ordered from smallest eigenvalue to the largest. Hence the projection of the gradient into the hessian can be written as follows

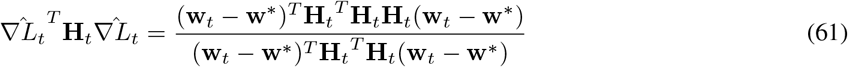

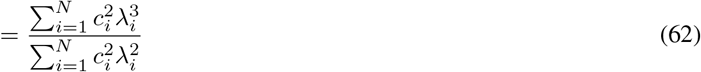

The hessian projection 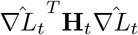 can range from the smallest eigenvalue *λ*_1_ (if **w**_*t*_ − **w** is in the direction of the 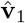) and *λ_N_* (if it is in the direction of 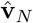). If the weight state **w**_*t*_ is chosen at random in weight space, **w**_*t*_ − **w**\* projects evenly onto the eigenvectors of the hessian. In that case, all the projection coefficients are equal *c_i_* = *c_j_* = *c* ∀*i, j* = 1,…, *N* and we get

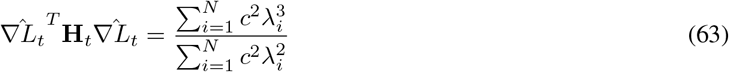

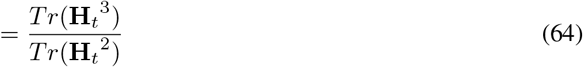

Note that in general, **w**_*t*_ is not chosen at random, it is determined by the weight trajectory from a particular learning rule. For example, if the learning rule is steepest gradient descent, the weights follow a trajectory that descends the steepest descent direction first. In that case, initially **w** _*t*_ − **w** is expected to project evenly onto all the eigenvectors, throughout learning **w**_*t*_ − **w** skews onto the directions with smaller curvature.

This gives us an explicit expression for the expected projection of the gradient ∇*L_t_*. What about the online gradient 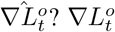 is an approximation of the task gradient ∇*L_t_* based on the available window of time as in eq (7).

In one extreme case, the online gradient is an exact approximation of the task gradient 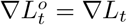. In that case, the hessian projections are equal. The less correlated the online gradient and the task gradient are, the smaller we expect this projection to be. Indeed, the only way the projection would increase is if the online gradient projected more strongly onto the directions of large curvature. Yet that would imply knowledge of the hessian of the task loss which is unlikely. Hence, we expect

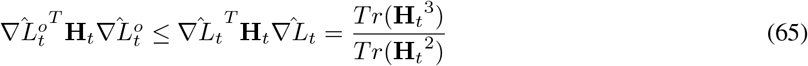

We can also bound the online gradient projection from the other side. Indeed, the other extreme case occurs when the online gradient is completely uncorrelated with the hessian. In that case, it projects evenly onto all the eigenvectors and 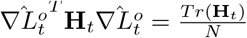. Combining these two results we obtain the following bound

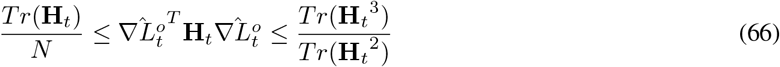

We have shown this bound is reasonable near a local minimum, where the task loss landscape can be approximated by a quadratic function. In general, the task loss will can be non-linear and non-convex. Hence, it might have multiple local minima, maxima and saddle points. Let **w**\* be a local minimum in the neighborhood of **w**. In this case, the hessian is no longer constant throughout weight space. We can write the gradient of the task loss at **w**_*t*_ as follows

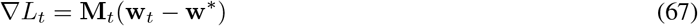

where

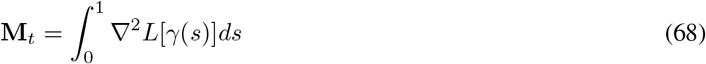

and *γ*(*s*) = *s***w**\* + (1 − *s*)**w**_*t*_, *s* ∈ [0, 1] is the parametrization of the straight line connecting **w**_*t*_ and **w**\*. The vector ∇*L_t_* can be decomposed in the directions of the eigenvectors of the hessian 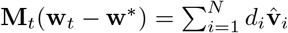. Hence the hessian projection of the gradient is

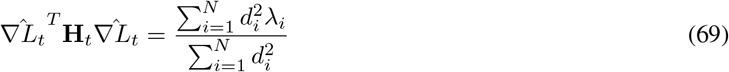

We consider two cases. First, close to steady state, when ||**w**_*t*_ − **w**\*|| is small, then **M**_*t*_ ≈ **H**_*t*_ and *d_i_* = *c_i_λ_i_* and the argument from the quadratic task loss applies. The bounds in (66) still hold.

Second, as ||**w**_*t*_ − **w**\*|| grows, further away from steady state, **M**_*t*_ is less similar than **H**_*t*_. If **M**_*t*_ and **H**_*t*_ were completely independent, then the projection of the gradient onto the different eigenvectors of the hessian is approximatively even and

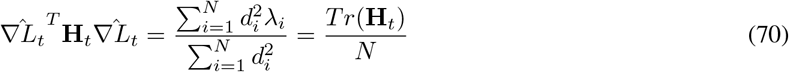

and the bounds in (66) are still valid.

Hence, for both linear and non-linear task loss functions and both at steady state and far from steady state the bounds in eq (16) are a good assumption. We make two further remarks. First, the better the online gradient approximation is (i.e. larger *γ°*), the closer the hessian projection will be to the upper bound. Second, at the beginning of learning, when the task loss overall decreases, we do not expect 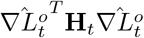 to be equal to the lower bound for long. If it were, the change in weights would be uncorrelated with the hessian and would behave like random error.

### Gradient and Hessian derivations

Both the gradient of the task loss and online loss have similar forms

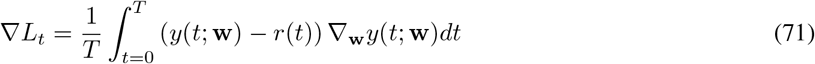

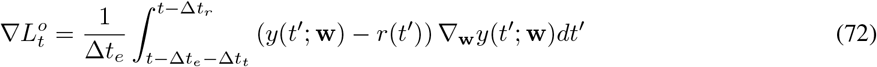

The gradients depend on the gradient of the plant output ∇_**w**_*y*(*t*; **w**). The output of the system depends on the network weights **w** through the plant input **u**.

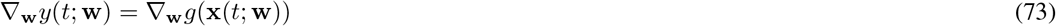

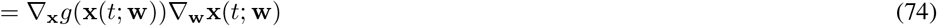

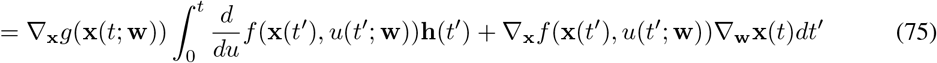

Hence the gradient of the task loss is

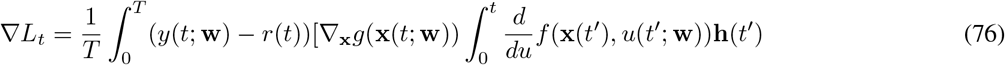

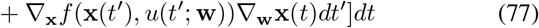

and the online gradient is

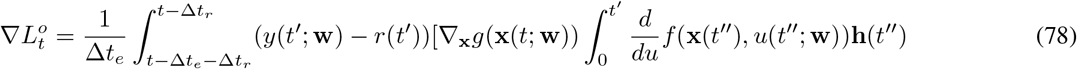

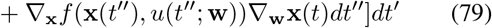

The hessian

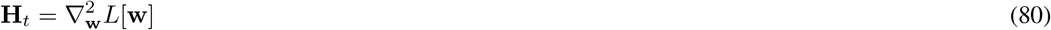

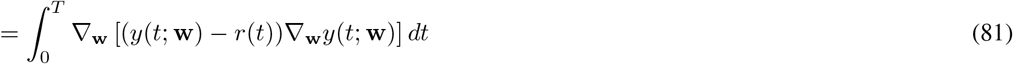

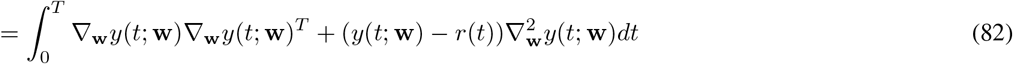

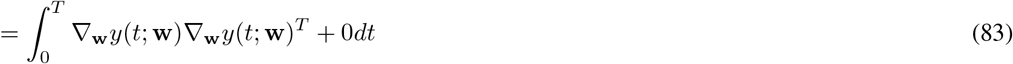

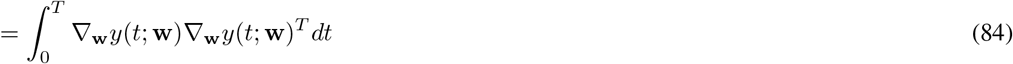

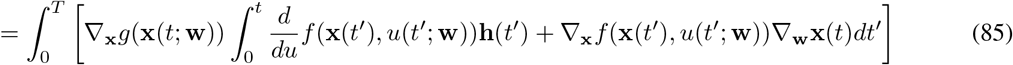

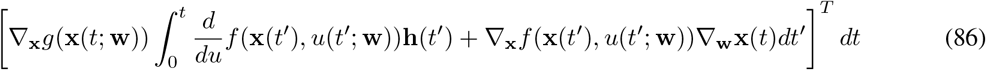

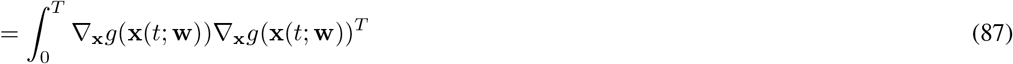

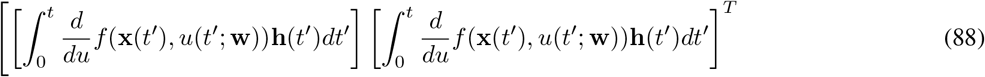

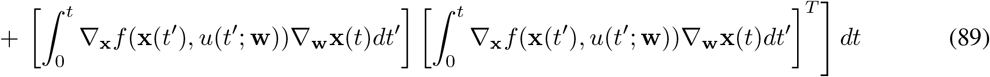

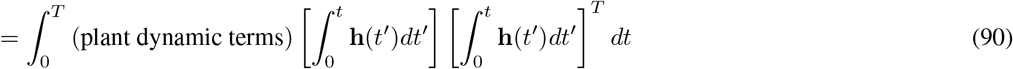

### Scaling of hessian projection terms

Lets define the matrix with the granule cell component of the Hessian matrix

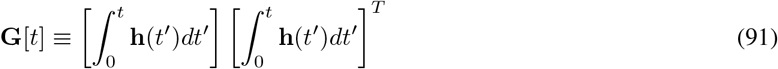

We assume the effect of the network expansion on the Hessian is mainly due to the change in the granule cell representation **h** and the matrix **G** and not the change in plant dynamics.

We found that the projection of the online gradient onto the Hessian is bound by *Tr*(**H**_*t*_)/*N* and *Tr*(**H**_*t*_^3^)/*Tr*(**H**_*t*_^2^) as in eq (16). Without knowing the plant dynamics we cannot calculate directly the projections in the initial and expanded network or the Hessian but we can consider how the quantities change relatively with the expansion.

We find that the trace increases linearly

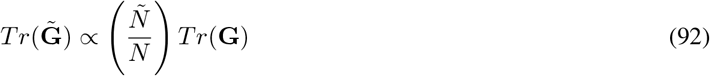

It follows that the average eigenvalue remains constant with the expansion

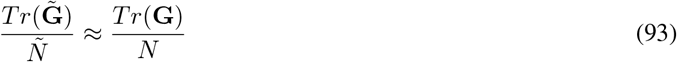

Furthermore, as the input expansion increases, the spread of the eigenvalues increases at most linearly.

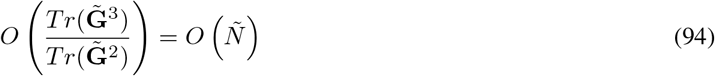

As we assume the effect of the network expansion on the Hessian is mainly due to the change in the granule cell representation **h** and the matrix **G** and not the change in plant dynamics. Then, the same scaling applies to the hessian terms.

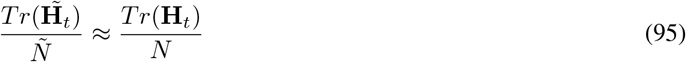

and

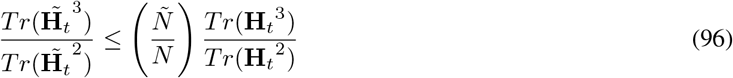

Using the bounds in eq (16), we get the following bounds for the hessian projection in the lager network.

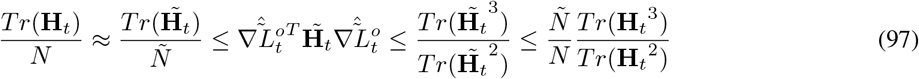

Hence, the projection of the online gradient onto the Hessian scales at most linearly with the network expansion.

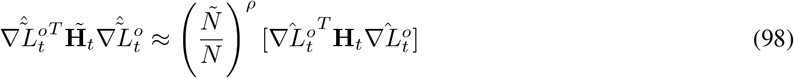

with 0 ≤ *ρ* ≤ 1 is the coefficient determining the scaling of the hessian projection.

We find that the Hessian projection 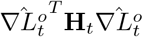 depends on 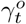. Indeed, the more correlated the online gradient is with the task gradient the more heavily it projects onto the directions of larger curvature. Similarly, *ρ* mainly depends on 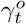. From simulations, we find the following fit

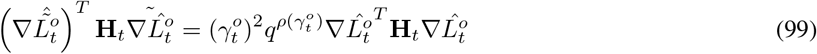

with

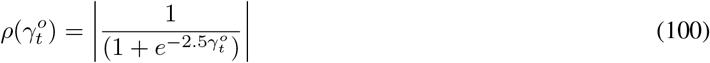

Hence, the less correlated the online gradient is with the task gradient (smaller 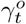, the slower it scales with *q*. Figure 1D shows the fit.

This has an effect on the scaling of learning performance as a function of *γ°*. The less correlated the online gradient is with the task gradient, the more beneficial the input expansion is for learning (Figure 2). We observe that as *γ°* increases, the learning speed increases slower with *q*. For large *γ°*, the learning speed initially increases, until it reaches a maximal value, beyond which the learning speed decreases. The larger *γ°* is, the larger the local task difficulty is. Indeed, near steady state, the effect of the curvature in the weight update is diminished when the change in weights is less correlated with the task gradient.

### Notes on simulations

We perform simulations of learning in the motor control system to test our theoretical results. We have three sources of variability in our simulations.

- from the reference trajectory *r*(*t*). The reference trajectory for a given simulation is drawn from a family of random reference trajectories as described in the methods. The variation comes from the selection of the phase shifts *ϕ_k_*

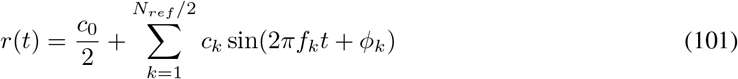

with

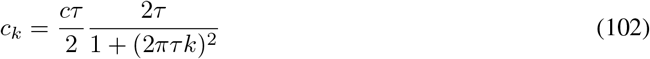

*r*(*t*) influences the task loss *L*[**w**]. We expect relatively little variation in the geometry of the task loss as a function of reference trajectory. More importantly, we expect the effect of the network expansion to be more significant than the variation from *r*(*t*). Indeed, each reference trajectory is chosen from the same family of trajectories (fourier series with the same cut-off frequency).
- from the weight initialization of both the input weights *Z* and the output weights **w**. This weights are initialised for each simulation according to the methods for the smallest network and then expanded for the larger networks. The value of *Z* affects directly the task loss *L* as it determines the mapping from the mossy fibre inputs to the granule cell layer activity *h*. Note that *Z* are constant during the whole trajectory as learning only changes the output weights. This value of the initial output weights **w**_0_ determines the initial point in the loss landscape, which determines both the initial task loss value *L*[**w**_0_] and the geometry around that point. We know that the optimal *γ*\* depends strongly on the geometry (slope and curvature) at the weight point. Hence this initialization will have a big effect on the increase of learning speed and decrease of steady state loss with respect to learning step *γ*. Furthermore, it will also determine the scaling of the learning parameters (gradient hessian projections) on the expansion ratio. The qualitative evolution for each weight initialization will be the same but the exact values will vary.
- from the learning rule error 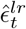 in the weight update rule (5).

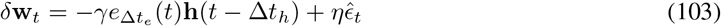

the vector 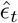 is drawn from a random distribution (see methods). We are interested in the expected learning speed and steady state loss over the learning rule error 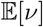 and 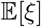. Hence we should train multiple times starting with the same initializations but with different learning rule error during training. This will lead to different weight trajectories and task loss during learning. We expect some variability in the learning speed and the steady state loss. The variability should be weaker than the change in learning performance from the change in expansion ratio.

**Figure 1:**
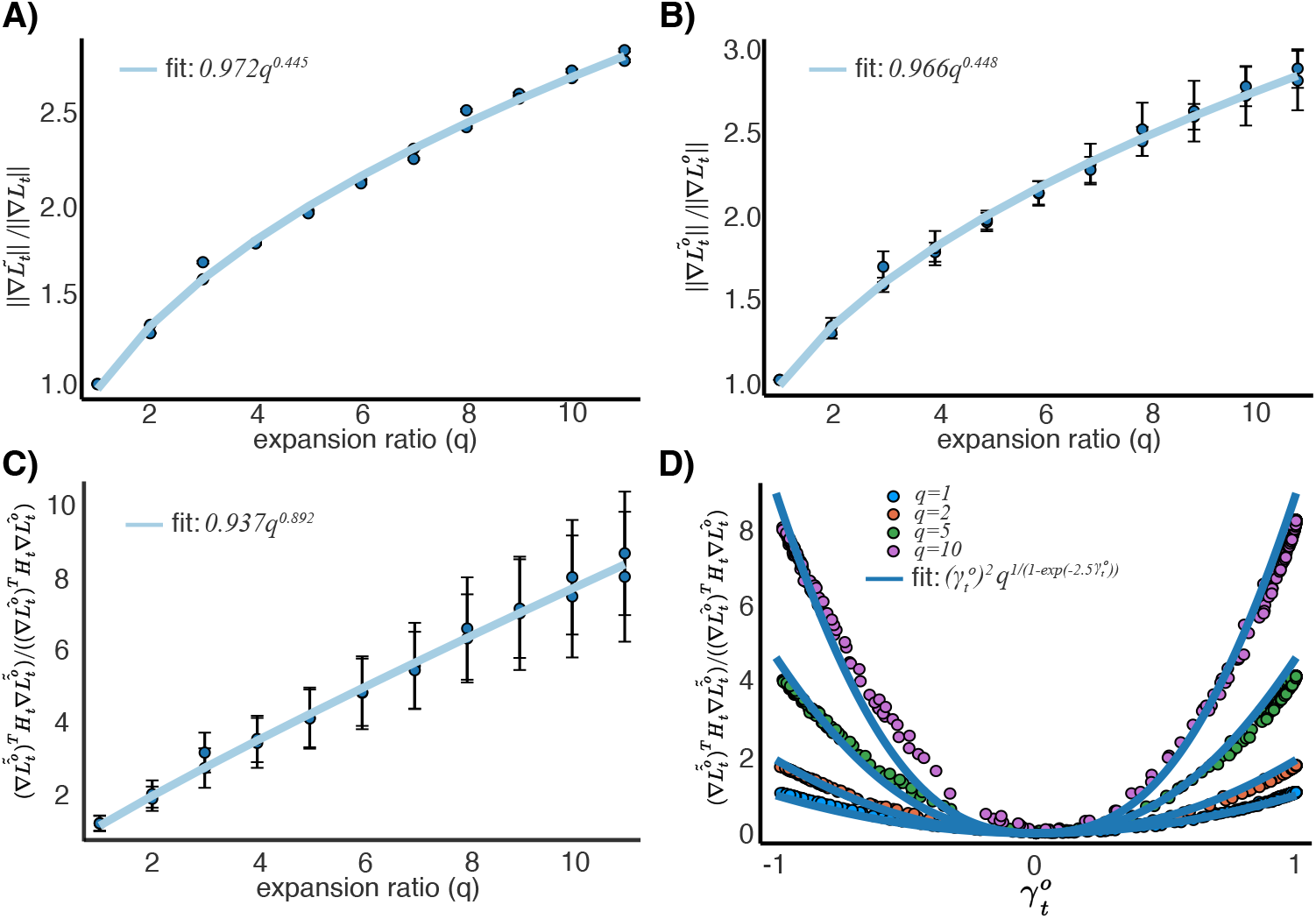
Scaling of learning parameters. **A)**, **B)**, **C)** We plot three different learning parameters for a range of expansion ratios *q* = 1,…, 11, for two different simulations with different weight initialisations and reference trajectories. We fit the data points with a power curve *y* = *ax^b^*. We recover the scaling parameters found in the results. **D)** We plot the normalised Hessian projection as a function of 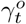 for different expansion ratios. Each color dot represents a different *q*. We plot the fit given by equations (99) and (100)

**Figure 2:**
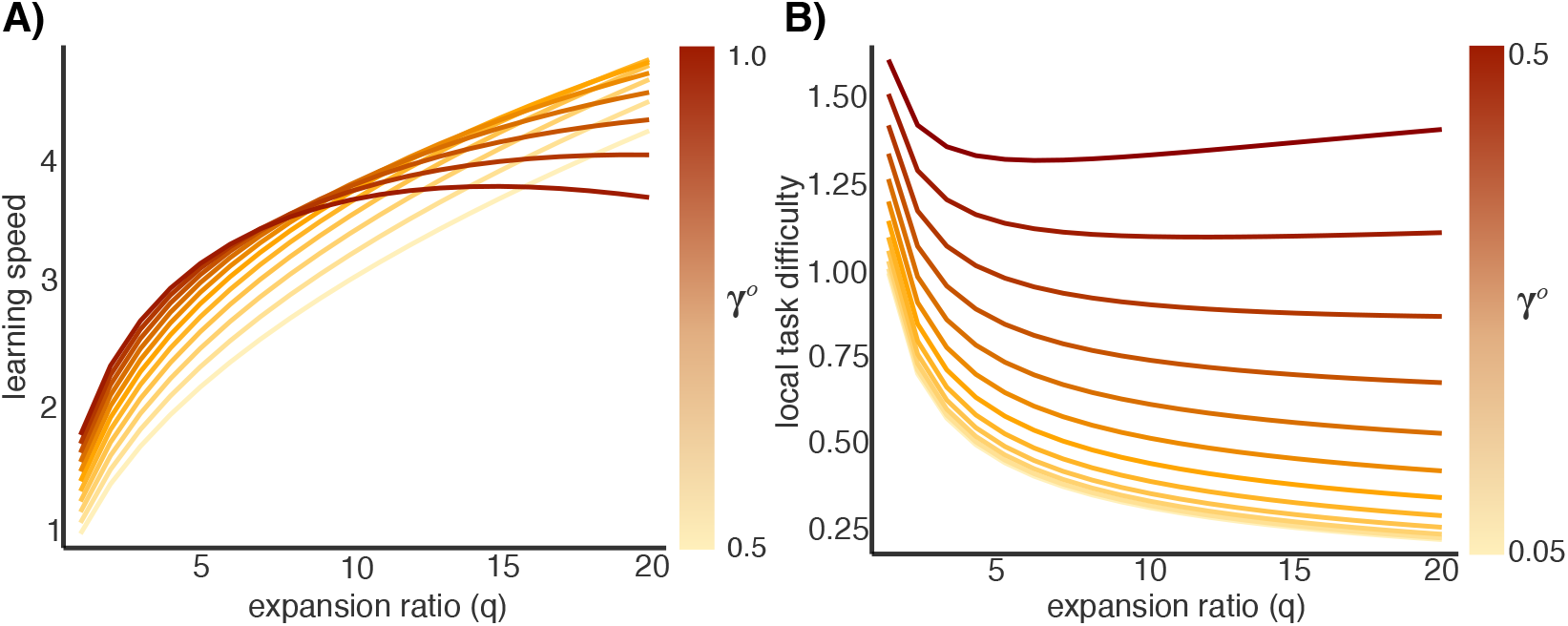
Effect of *γ°* on learning speed **A)** and local task difficulty **B)**. We plot the learning performance parameter as a function of the expansion ratio *q*, for different *γ°*. The color of the line represents *γ°*.

In Figure 2C and D each dot is the mean over different simulations with different *r*(*t*) and weight initialisations. This guarantees that our results are not dependent on the choice of reference trajectory or weight initialisation. It just requires being in a regime in which the system is learning (decreasing the task loss). The scatter plot B) showing the trade-off between learning speed and steady state loss is plotted for a single *r*(*t*) and weight initialisation but for 20 different training instances. As mentioned above the weight initialisation can have a strong effect on the geometry of the task loss and hence on the dependence of *ν* and *ξ* on *γ* and the scaling with respect to *q*. The results of the trade-off and will still hold but the exact factors will vary.

**Figure 3:**
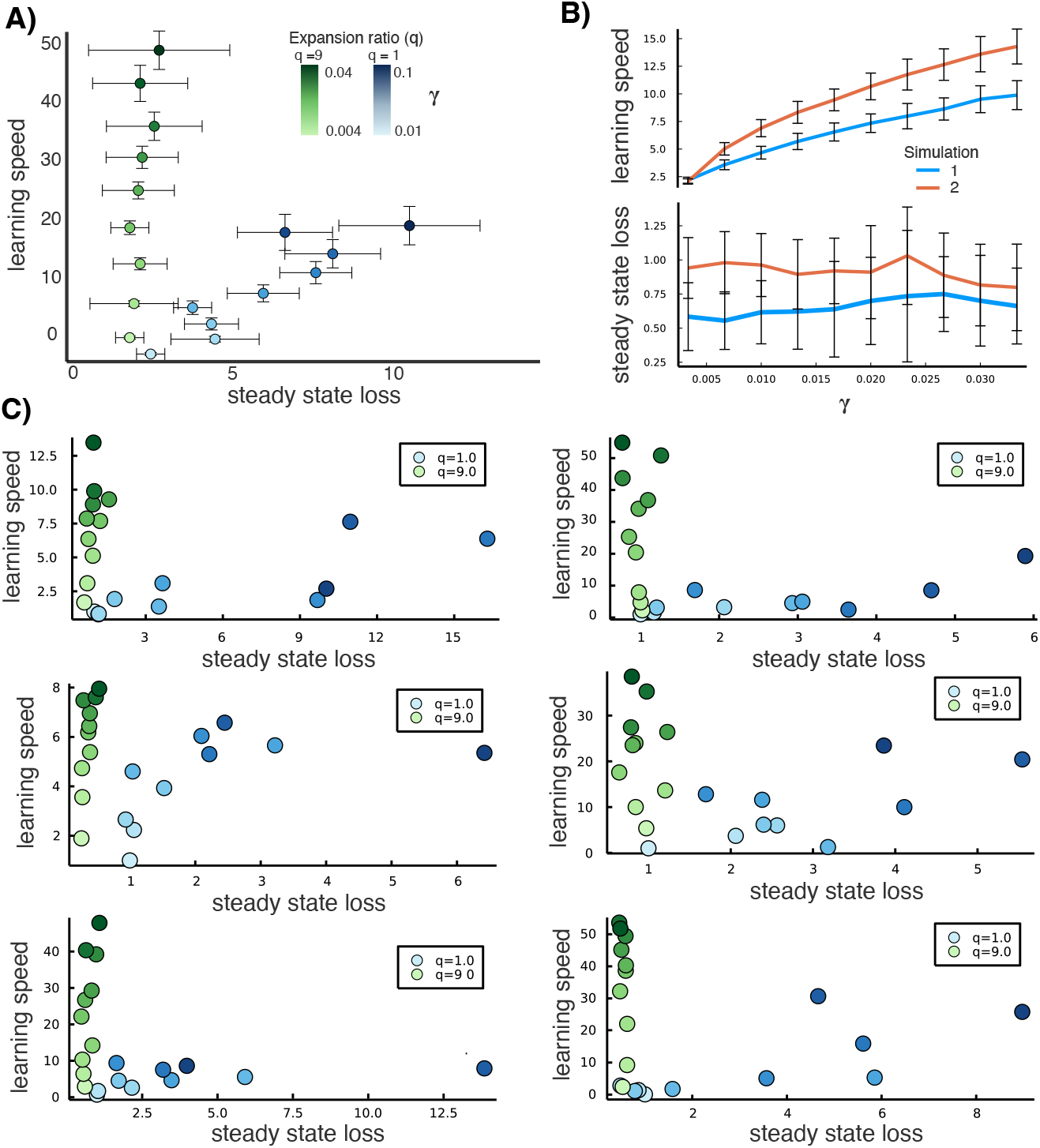
Effect of variability in simulations. **A)** Trade-off between steady state loss and learning speed averaged over 17 different reference trajectories and weight initilizations. Same data as in Figure 2C and D. The variability is high. **B)** Learning speed and steady state loss as a function of *γ* for two different simulations. For both simulations the trend in the learning variables is the same but coefficients are different. **C)** Trade-off of steady state loss and learning speed for a subset of 6 different simulations used for **A)**. For each different simulation the trade-off is visible but has different values.

## Notes

### Competing Interest Statement

The authors have declared no competing interest.

### Summary of Updates

This version has been revised to update the overall language of the manuscript without changing the results.

## References

1 James S. Albus. A theory of cerebellar function. Mathematical Biosciences, 10(1-2):25–61, 1971.

2 David Marr. A theory of cerebellar cortex. The Journal of Physiology, 202(2):437–470, jun 1969.

3 Ashok Litwin-Kumar, Kameron Decker Harris, Richard Axel, Haim Sompolinsky, and L. F. Abbott. Optimal Degrees of Synaptic Connectivity. Neuron, 2017.

4 N. Alex Cayco-Gajic, Claudia Clopath, and R. Angus Silver. Sparse synaptic connectivity is required for decorrelation and pattern separation in feedforward networks. Nature Communications, 8(1), dec 2017.

5 John C Eccles. The Cerebellum as a Computer: Patterns in Space and Time. J. Phygiol, 229:1–32, 1973.

6 Terence D. Sanger, Okito Yamashita, and Mitsuo Kawato. Expansion coding and computation in the cerebellum: 50 years after the Marr–Albus codon theory. The Journal of Physiology, 598(5):913–928, mar 2020.

7 N Alex Cayco-Gajic and R Angus Silver. Re-evaluating Circuit Mechanisms Underlying Pattern Separation. Neuron, 101(4):584–602, 2019.

8 Guy Billings, Eugenio Piasini, Andrea Lorincz, Zoltan Nusser, and R. Angus Silver. Network Structure within the Cerebellar Input Layer Enables Lossless Sparse Encoding. Neuron, 83(4):960–974, aug 2014.

9 Baktash Babadi and Haim Sompolinsky. Sparseness and expansion in sensory representations. Neuron, 83:1213–1226, 9 2014.

10 Anton Spanne and Henrik Jörntell. Questioning the role of sparse coding in the brain. Trends in Neurosciences, 38:417–427, 7 2015.

11 Mitsuo Kawato, Shogo Ohmae, Huu Hoang, and Terry Sanger. 50 Years Since the Marr, Ito, and Albus Models of the Cerebellum, jun 2020.

12 Laura D. Knogler, Daniil A. Markov, Elena I. Dragomir, Vilim Štih, and Ruben Portugues. Sensorimotor Representations in Cerebellar Granule Cells in Larval Zebrafish Are Dense, Spatially Organized, and Non-temporally Patterned. Current Biology, 27(9):1288–1302, may 2017.

13 Mark J. Wagner, Tony Hyun Kim, Joan Savall, Mark J. Schnitzer, and Liqun Luo. Cerebellar granule cells encode the expectation of reward. Nature, 544(7648):96–100, apr 2017.

14 Andrea Giovannucci, Aleksandra Badura, Ben Deverett, Farzaneh Najafi, Talmo D. Pereira, Zhenyu Gao, Ilker Ozden, Alexander D. Kloth, Eftychios Pnevmatikakis, Liam Paninski, Chris I. De Zeeuw, Javier F. Medina, and Samuel S.H. Wang. Cerebellar granule cells acquire a widespread predictive feedback signal during motor learning. Nature Neuroscience, 20(5):727–734, may 2017.

15 Aleksandra Badura and Chris I. De Zeeuw. Cerebellar Granule Cells: Dense, Rich and Evolving Representations, jun 2017.

16 Jesse I. Gilmer and Abigail L. Person. Theoretically Sparse, Empirically Dense: New Views on Cerebellar Granule Cells, dec 2018.

17 Susanne M. Morton and Amy J. Bastian. Mechanisms of cerebellar gait ataxia. Cerebellum, 6(1):79–86, 2007.

18 Maurice A. Smith, Jason Brandt, and Reza Shadmehr. Motor Disorder in Huntington’s Disease Begins as a Dysfunction in Error Feedback Control. Nature, 403(6769):544, feb 2000.

19 Sarah E. Criscimagna-Hemminger, Amy J. Bastian, and Reza Shadmehr. Size of error affects cerebellar contributions to motor learning. Journal of Neurophysiology, 103:2275–2284, 4 2010.

20 Tycho M. Hoogland, Jornt R. De Gruijl, Laurens Witter, Cathrin B. Canto, and Chris I. De Zeeuw. Role of synchronous activation of cerebellar purkinje cell ensembles in multi-joint movement control. Current biology : CB, 25:1157–1165, 5 2015.

21 Ana S. Machado, Dana M. Darmohray, João Fayad, Hugo G. Marques, and Megan R. Carey. A quantitative framework for whole-body coordination reveals specific deficits in freely walking ataxic mice. eLife, 4, 10 2015.

22 Mitsuo Kawato, Kazunori Furukawa, and R. Suzuki. A hierarchical neural-network model for control and learning of voluntary movement. Biological Cybernetics, 57(3):169–185, oct 1987.

23 Mitsuo Kawato and Hiroaki Gomi. A computational model of four regions of the cerebellum based on feedback-error learning. Biological Cybernetics, 68(2):95–103, dec 1992.

24 Sergio Oscar Verduzco-Flores and Erik De Schutter. Self-configuring feedback loops for sensorimotor control. eLife, 11, 11 2022.

25 Hiroaki Gomi and Mitsuo Kawato. Equilibrium-point control hypothesis examined by measured arm stiffness during multijoint movement. Science, 272(5258):117–120, apr 1996.

26 Brendan D. Cameron, Cristina de la Malla, and Joan López-Moliner. The role of differential delays in integrating transient visual and proprioceptive information. Frontiers in Psychology, 5:50, 2014.

27 R C Miall, D J Weir, D M Wolpert, and J F Stein. Is the Cerebellum a Smith Predictor? Journal of Motor Behavior, 25(3):203–216, 1993.

28 R. C. Miall and Daniel M. Wolpert. Forward models for physiological motor control. Neural Networks, 9:1265–1279, 1996.

29 Jeffrey A. Saunders and David C. Knill. Visual feedback control of hand movements. Journal of Neuroscience, 24:3223–3234, 3 2004.

30 Sergio Verduzco-Flores, William Dorrell, and Erik De Schutter. A differential hebbian framework for biologically-plausible motor control. Neural Networks, 150:237–258, 6 2022.

31 Daniel M. Wolpert, R. Chris Miall, and Mitsuo Kawato. Internal models in the cerebellum. Trends in Cognitive Sciences, 2(9):338–347, sep 1998.

32 Chris I. De Zeeuw, Stephen G. Lisberger, and Jennifer L. Raymond. Diversity and dynamism in the cerebellum. Nature Neuroscience, pages 1–8, 12 2020.

33 A. J. Bastian, T. A. Martin, J. G. Keating, and W. T. Thach. Cerebellar ataxia: Abnormal control of interaction torques across multiple joints. Journal of Neurophysiology, 76(1):492–509, 1996.

34 Susanne M. Morton and Amy J. Bastian. Cerebellar contributions to locomotor adaptations during splitbelt treadmill walking. Journal of Neuroscience, 26:9107–9116, 9 2006.

35 K. Rabe, O. Livne, E. R. Gizewski, V. Aurich, A. Beck, D. Timmann, and O. Donchin. Adaptation to visuomotor rotation and force field perturbation is correlated to different brain areas in patients with cerebellar degeneration. Journal of Neurophysiology, 101:1961–1971, 4 2009.

36 Maurice A. Smith and Reza Shadmehr. Intact ability to learn internal models of arm dynamics in huntington’s disease but not cerebellar degeneration. Journal of Neurophysiology, 93:2809–2821, 5 2005.

37 Nasir H Bhanpuri, Allison M Okamura, and Amy J Bastian. Predicting and correcting ataxia using a model of cerebellar function. Brain : a journal of neurology, 137(Pt 7):1931–44, jul 2014.

38 David J. Herzfeld, Yoshiko Kojima, Robijanto Soetedjo, and Reza Shadmehr. Encoding of error and learning to correct that error by the Purkinje cells of the cerebellum. Nature Neuroscience, 21(5):736–743, may 2018.

39 Shigeru Kitazawa, Tatsuya Kimura, and Ping Bo Yin. Cerebellar complex spikes encode both destinations and errors in arm movements. Nature 1998 392:6675, 392(6675):494–497, apr 1998.

40 Martha L. Streng, Laurentiu S. Popa, and Timothy J. Ebner. Climbing fibers control purkinje cell representations of behavior. Journal of Neuroscience, 37(8):1997–2009, feb 2017.

41 Yan Yang and Stephen G. Lisberger. Modulation of complex-spike duration and probability during cerebellar motor learning in visually guided smooth-pursuit eye movements of monkeys. eNeuro, 4, 6 2017.

42 Yunliang Zang and Erik De Schutter. Climbing fibers provide graded error signals in cerebellar learning. Frontiers in Systems Neuroscience, 13:46, 9 2019.

43 Javier F. Medina and Stephen G. Lisberger. Links from complex spikes to local plasticity and motor learning in the cerebellum of awake-behaving monkeys. Nature Neuroscience, 11:1185–1192, 10 2008.

44 Yan Yang and Stephen G. Lisberger. Purkinje-cell plasticity and cerebellar motor learning are graded by complex-spike duration. Nature, 510:529–532, 2014.

45 Aparna Suvrathan, Hannah L. Payne, and Jennifer L. Raymond. Timing Rules for Synaptic Plasticity Matched to Behavioral Function. Neuron, 92(5):959–967, dec 2016.

46 Patrick Safo and Wade G. Regehr. Timing dependence of the induction of cerebellar ltd. Neuropharmacology, 54:213–218, 1 2008.

47 Dhruva V. Raman and Timothy O’Leary. Frozen algorithms: how the brain’s wiring facilitates learning. Current Opinion in Neurobiology, 67:207–214, 4 2021.

48 B Widrow and ME Hoff. Adaptive switching circuits. 1960.

49 Kenji Yamamoto, Yasushi Kobayashi, Aya Takemura, Kenji Kawano, and Mitsuo Kawato. Computational studies on acquisition and adaptation of ocular following responses based on cerebellar synaptic plasticity. Journal of Neurophysiology, 87:1554–1571, 2002.

50 Dhruva V. Raman and Timothy O’leary. Optimal plasticity for memory maintenance during ongoing synaptic change. eLife, 10, sep 2021.

51 Dhruva Venkita Raman, Adriana Perez Rotondo, and Timothy O’Leary. Fundamental bounds on learning performance in neural circuits. Proceedings of the National Academy of Sciences of the United States of America, 116(21):10537–10546, may 2019.

52 Marjorie Xie, Samuel Muscinelli, Kameron Decker Harris, and Ashok Litwin-Kumar. Task-dependent optimal representations for cerebellar learning. bioRxiv, 2022.

53 T. Tyrrell and D. Willshaw. Cerebellar cortex: its simulation and the relevance of marr’s theory. Philosophical transactions of the Royal Society of London. Series B, Biological sciences, 336:239–257, 5 1992.

54 Toyoshi Torioka. Pattern separability and the effect of the number of connections in a random neural net with inhibitory connections. Biol. Cybernetics, 31:27–35, 1978.

55 N. Schweighofer, K. Doya, and F. Lay. Unsupervised learning of granule cell sparse codes enhances cerebellar adaptive control. Neuroscience, 103:35–50, 2 2001.

56 Tri M Nguyen, Logan A Thomas, Jeff L Rhoades, Ilaria Ricchi, Xintong Cindy, Arlo Sheridan, David G C Hildebrand, Jan Funke, Wade G Regehr, and Allen Lee. Structured connectivity in the cerebellum enables noise-resilient pattern separation. 2021.

57 Asha Vijayan and Shyam Diwakar. A cerebellum inspired spiking neural network as a multi-model for pattern classification and robotic trajectory prediction. Frontiers in Neuroscience, 16:2004, 11 2022.

58 Henrik Jörntell and Carl Fredrik Ekerot. Properties of somatosensory synaptic integration in cerebellar granule cells in vivo. Journal of Neuroscience, 26:11786–11797, 11 2006.

59 Jesse I. Gilmer, Michael A. Farries, Zachary Kilpatrick, Ioannis Delis, and Abigail L. Person. An emergent temporal basis set robustly supports cerebellar time-series learning. bioRxiv, page 2022.01.06.475265, 1 2022.

60 Laurens Witter and Chris I. De Zeeuw. In vivo differences in inputs and spiking between neurons in lobules vi/vii of neocerebellum and lobule x of archaeocerebellum. Cerebellum, 14:506–515, 10 2015.

61 Isabelle Straub, Laurens Witter, Abdelmoneim Eshra, Miriam Hoidis, Niklas Byczkowicz, Sebastian Maas, Igor Delvendahl, Kevin Dorgans, Elise Savier, Ingo Bechmann, Martin Krueger, Philippe Isope, and Stefan Hallermann. Gradients in the mammalian cerebellar cortex enable fourier-like transformation and improve storing capacity. eLife, 9, 2 2020.

62 Frank Van Overwalle, Mario Manto, Zaira Cattaneo, Silvia Clausi, Chiara Ferrari, John D.E. Gabrieli, Xavier Guell, Elien Heleven, Michela Lupo, Qianying Ma, Marco Michelutti, Giusy Olivito, Min Pu, Laura C. Rice, Jeremy D. Schmahmann, Libera Siciliano, Arseny A. Sokolov, Catherine J. Stoodley, Kim van Dun, Larry Vandervert, and Maria Leggio. Consensus paper: Cerebellum and social cognition. Cerebellum, 19:833–868, 12 2020.

63 Adele Diamond. Close interrelation of motor development and cognitive development and of the cerebellum and prefrontal cortex. Child development, 71:44–56, 2000.

64 Hanne Baillieux, Hyo Jung De Smet, Philippe F. Paquier, Peter P. De Deyn, and Peter Mariën. Cerebellar neurocognition: insights into the bottom of the brain. Clinical neurology and neurosurgery, 110:763–773, 8 2008.

65 M Küper, A Dimitrova, M Thürling, S Maderwald, J Roths, H G Elles, E R Gizewski, M E Ladd, J Diedrichsen, and D Timmann. Evidence for a motor and a non-motor domain in the human dentate nucleus — an fmri study. 2011.

66 Marco Molinari, Francesca R Chiricozzi, Silvia Clausi, Anna Maria Tedesco, Mariagrazia De Lisa, and Maria G Leggio. Cerebellum and detection of sequences, from perception to cognition. Cerebellum, pages 611–615, 2008.

67 Masao Ito. Movement and thought: identical control mechanisms by the cerebellum. Trends in Neurosciences, 16:448–450, 11 1993.

68 John Porrill, Paul Dean, and Sean R. Anderson. Adaptive filters and internal models: Multilevel description of cerebellar function. Neural Networks, 47:134–149, 11 2013.

69 Michael I. Jordan and David E. Rumelhart. Forward models: Supervised learning with a distal teacher. Cognitive Science, 16:307–354, 7 1992.

70 Michael Paulin. A Kalman Filter Theory of the Cerebellum. 1989.

71 Alberto Antonietti, Claudia Casellato, Jesús A. Garrido, Niceto R. Luque, Francisco Naveros, Eduardo Ros, Egidio D’Angelo, and Alessandra Pedrocchi. Spiking neural network with distributed plasticity reproduces cerebellar learning in eye blink conditioning paradigms. IEEE Transactions on Biomedical Engineering, 63:210–219, 1 2016.

72 Asha Vijayan, Vivek Gopan, Bipin Nair, and Shyam Diwakar. Comparing robotic control using a spiking model of cerebellar network and a gain adapting forward-inverse model. volume 2017-Janua, pages 566–570. Institute of Electrical and Electronics Engineers Inc., 11 2017.

73 Claudia Casellato, Alberto Antonietti, Jesus A. Garrido, Richard R. Carrillo, Niceto R. Luque, Eduardo Ros, Alessandra Pedrocchi, and Egidio D’Angelo. Adaptive Robotic Control Driven by a Versatile Spiking Cerebellar Network. PLoS ONE, 9(11):e112265, nov 2014.

74 Richard R. Carrillo, Eduardo Ros, Christian Boucheny, and Olivier J.M.D. Coenen. A real-time spiking cerebellum model for learning robot control. BioSystems, 94(1-2):18–27, oct 2008.

75 Stéphane Dieudonné. Submillisecond kinetics and low efficacy of parallel fibre-golgi cell synaptic currents in the rat cerebellum. The Journal of Physiology, 510:845, 8 1998.

76 Henrik Jörntell, Fredrik Bengtsson, Martijn Schonewille, and Chris I. De Zeeuw. Cerebellar molecular layer interneurons – computational properties and roles in learning. Trends in Neurosciences, 33:524–532, 11 2010.

77 James M. Bower. The organization of cerebellar cortical circuitry revisited. Annals of the New York Academy of Sciences, 978:135–155, 12 2002.

78 Rodolfo R. Llinás, Kerry D. Walton, and Eric J. Lang. Cerebellum. The Synaptic Organization of the Brain, 1 2004.

79 Elizabeth A. Fleming, Michael R. Tadross, and Court Hull. Local synaptic inhibition mediates cerebellar pattern separation necessary for learned sensorimotor associations. bioRxiv, page 2022.05.20.492839, 5 2022.

80 Masao Ito and Masanobu Kano. Long-lasting depression of parallel fiber-Purkinje cell transmission induced by conjunctive stimulation of parallel fibers and climbing fibers in the cerebellar cortex. Neuroscience Letters, 33(3):253–258, dec 1982.

81 T. Hirano. Depression and potentiation of the synaptic transmission between a granule cell and a Purkinje cell in rat cerebellar culture. Neuroscience Letters, 119(2):141–144, nov 1990.

82 Megan R. Carey. Synaptic mechanisms of sensorimotor learning in the cerebellum. Current Opinion in Neurobiology, 21:609–615, 8 2011.

83 N. Tatiana Silva, Jorge Ramírez-Buriticá, Dominique L. Pritchett, and Megan R. Carey. Neural instructive signals for associative cerebellar learning. bioRxiv, page 2022.04.18.488634, 4 2022.

84 Jennifer L. Raymond and Javier F. Medina. Computational Principles of Supervised Learning in the Cerebellum. Annual Review of Neuroscience, 41(1):233–253, jul 2018.

85 Chris I. De Zeeuw, Stephen G. Lisberger, and Jennifer L. Raymond. Diversity and dynamism in the cerebellum. Nature Neuroscience, 24:160–167, 12 2020.

86 Stephen G. Lisberger. The rules of cerebellar learning: Around the ito hypothesis. Neuroscience, 462:175–190, 5 2021.

87 Zhenyu Gao, Boeke J. Van Beugen, and Chris I. De Zeeuw. Distributed synergistic plasticity and cerebellar learning, sep 2012.

